# Molecular basis of F-actin regulation and sarcomere assembly *via* myotilin

**DOI:** 10.1101/2020.09.25.310029

**Authors:** Julius Kostan, Miha Pavšič, Vid Puž, Thomas C. Schwarz, Friedel Drepper, Sibylle Molt, Melissa Ann Graewert, Claudia Schreiner, Sara Sajko, Peter F.M. van der Ven, Dmitri I. Svergun, Bettina Warscheid, Robert Konrat, Dieter O. Fürst, Brigita Lenarčič, Kristina Djinović-Carugo

**Affiliations:** Department of Structural and Computational Biology, Max Perutz Labs, University of Vienna, Campus Vienna Biocenter 5, A-1030 Vienna, Austria; Department of Biochemistry, Faculty of Chemistry and Chemical Technology, University of Ljubljana, Večna pot 5, SI-1000 Ljubljana, Slovenia; Biochemistry and Functional Proteomics, Institute of Biology II, Faculty of Biology, University of Freiburg, 79104, Freiburg, Germany; Signalling Research Centres BIOSS and CIBSS, University of Freiburg, 79104, Freiburg, Germany; Institute for Cell Biology, Department of Molecular Cell Biology, University of Bonn, Ulrich- Haberland-Str. 61a, 53121 Bonn, Germany; European Molecular Biology Laboratory, Hamburg Unit, c/o DESY, Notkestrasse 85, 22607 Hamburg, Germany; Department of Biochemistry, Molecular Biology and Structural Biology, Jozef Stefan Institute, Jamova 39, SI-1000 Ljubljana, Slovenia

**Author notes:** Correspondence and requests for materials should be addressed to KD-C and/or BL. These authors contributed equally to this work (Julius Kostan, Miha Pavšič).

**Keywords:** Sarcomeric Z-disc, F-actin binding, myotilin, α-Actinin, integrative structural model of F-actin/myotilin, competitive tropomyosin binding, Z-disc-assembly regulation

## Abstract

Sarcomeres, the basic contractile units of striated muscle cells, contain arrays of thin (actin) and thick (myosin) filaments that slide past each other during contraction. The Ig-like domain containing protein myotilin provides structural integrity to Z-discs - the boundaries between adjacent sarcomeres. Myotilin binds to Z-disc components, including F-actin and α-actinin-2, but the molecular mechanism of binding and implications of these interactions on Z-disc integrity are still elusive. We used a combination of small angle X-ray scattering, cross-linking mass spectrometry, biochemical and molecular biophysics approaches. We discovered that myotilin displays conformational ensembles in solution. We generated a structural model of the F-actin:myotilin complex that revealed how myotilin interacts with and stabilizes F-actin *via* its Ig-like domains and flanking regions. Mutant myotilin designed with impaired F-actin binding showed increased dynamics in cells. Structural analyses and competition assays uncovered that myotilin displaces tropomyosin from F-actin. Our findings suggest a novel role of myotilin as a co-organizer of Z-disc assembly and advance our mechanistic understanding of myotilin’s structural role in Z-discs.

**Significance Statement:** Sarcomeres are the primary structural and functional unit of striated muscles, conferring movement in all animals. The Z-disk is the boundary between adjacent sarcomeres, where actin filaments (F-actin) are anchored. Z-disc protein myotilin, is a scaffold protein, which provides structural integrity to the Z-disc by multiple interactions to its central components, including F-actin and α-actinin-2. Here we provide the structure of myotilin, revealing its structural plasticity in solution and the first integrative structural model of its complex with F-actin. We further show that myotilin displaces tropomyosin from F-actin, implying a novel role of myotilin in sarcomere biogenesis beyond being an interaction hub for Z-disk partners.

**Highlights:**

፧ Myotilin is structurally described as a dynamic ensemble
፧ Flanking regions enhance F-acting binding to tandem Ig domains
፧ Integrative structural model of myotilin bound to F-actin
፧ Myotilin displaces tropomyosin from F-actin, suggesting an organisational role in Z-disc

## Introduction

About 40% of the human body is comprised of skeletal muscle, whose contraction leads to locomotion [1]. The contractile machinery of cross-striated muscle cells is based on an impressive, almost crystalline array of thin (actin-based) and thick (myosin-based) filaments, arranged in repeating units, the sarcomeres. Stringent control of the precise layout of these filament systems is of utmost importance for efficient conversion of the force produced by the myosin-actin-interaction into contraction at the macroscopic level. Thin filaments are cross-linked in an antiparallel fashion at the Z-discs, the boundaries of adjacent sarcomeres, by multiple molecular interactions (Figure 1A). In fact, more than 50 proteins may be associated with mature Z-discs and they are regarded as one of the most complex macromolecular structures in biology. For decades the Z-disc was believed to play a specific role only in sustaining myofibril architecture. This view has changed dramatically within the last decade, with the Z-disc now also being recognized as a prominent hub for signalling, mechanosensing and mechanotransduction, with emerging roles in protein turnover and autophagy [2, 3]. In line with this, Z-disc proteins have recently been identified as a major phosphorylation hotspot, thereby directly modulating protein interactions and dynamics [4, 5].

**Figure 1.**
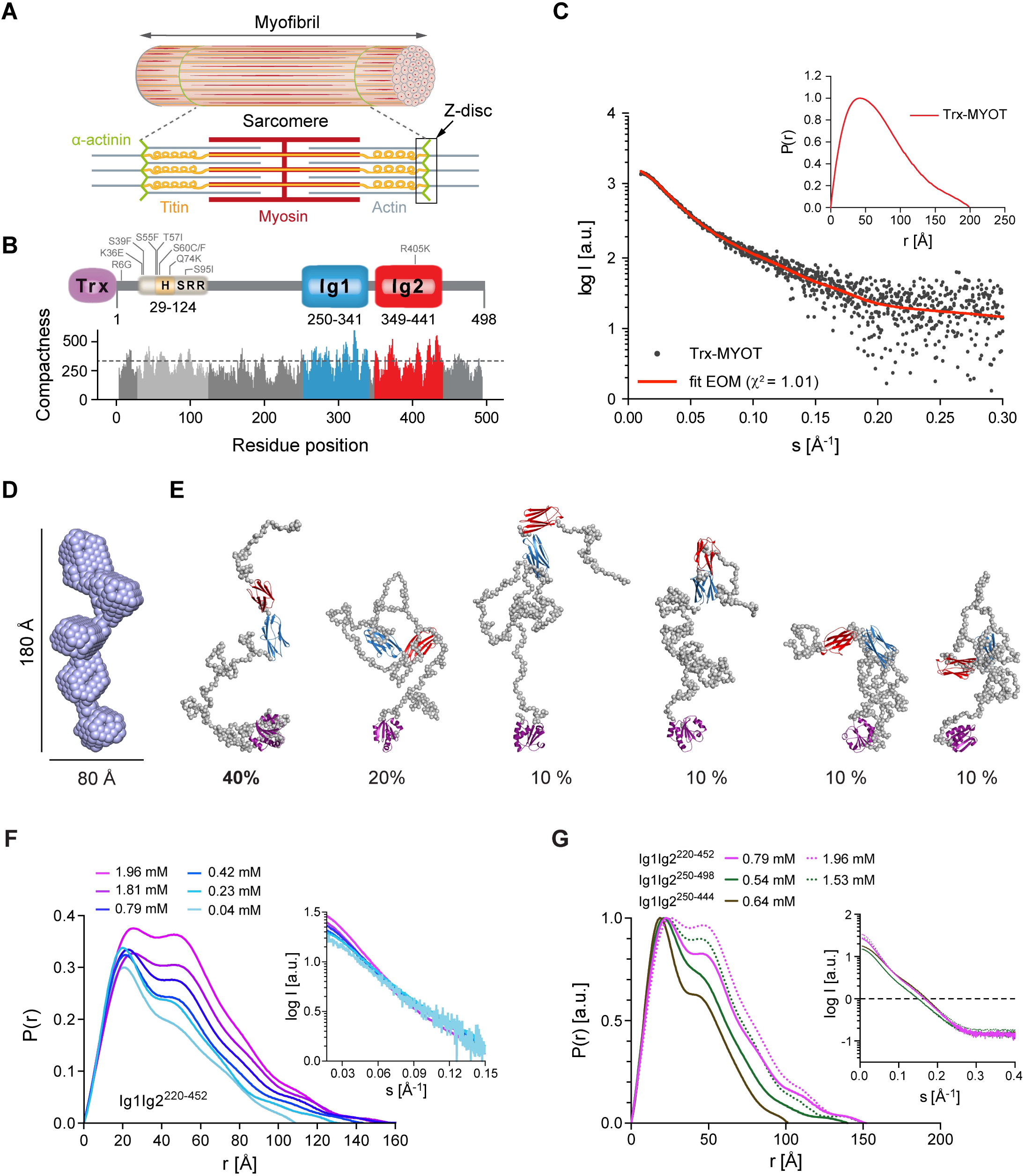
Myotilin displays conformational ensemble in solution. (**A**) Schematic representation of striated muscle sarcomere with its major thick (myosin-based) and thin (actin-based) filaments, titin and the Z-disc anchoring cross-linker α-actinin. Extending from one Z-disc to the next, sarcomere represents the fundamental unit of muscle contraction. (**B**) Schematic presentation of myotilin, and its predicted compactness as a function of residue position. The N-terminal part of myotilin contains the serine-rich region (SRR), which comprise hydrophobic residues stretch (H, yellow) and represents “mutational hotspot” of the protein (grey). Known disease-causing mutations are shown. C-terminal Ig domains 1 and 2, are coloured blue and red, respectively, followed by the C-terminal tail. Within this study, full-length myotilin construct fused to N-terminal thioredoxin (Trx, violet), was used. Large meta-structure derived compactness values indicate residue positions typically buried in the interior of the 3D structure, whereas small values are found for residues exposed to the solvent. Dashed line depicts the average compactness value (about 300) of the well-folded protein. Significantly smaller values (200) are found for structurally flexible proteins [23]. (**C**) Experimental SAXS scattering data of Trx-MYOT, with the fit of the EOM flexible modelling. Inset, *P*(*r*) *vs. r* plot for Trx-MYOT with the D_max_ □200 Å. (**D**) SAXS-based *ab initio* molecular envelope of Trx-MYOT. Most probable model is shown. (**E**) EOM models of Trx-MYOT showing flexible and intrinsically disordered N- and C-terminal regions (grey). The selected models are presented with the percentage contribution, estimated from the final population of EOM models. (**F**) *P(r) vs. r* plot for concentration series of Ig1Ig2^220-452^. Inset, respective concentration series with the corresponding SAXS profiles. (**G**) *P(r) vs. r* plot for dimeric Ig1Ig2^220-452^, Ig1Ig2^250-498^ and monomeric Ig1Ig2^220-444^. Inset, SAXS profiles of respective constructs. In order to compare various *P(r)* functions, *P(r)* was normalized to the peak height. See also **Figure S1 and Table S1**.

A striking ultrastructural feature of the Z-disc is its highly ordered, paracrystalline tetragonal arrangement [6]. This arrangement is governed by α-actinin-2, which cross-links the overlapping ends of thin filaments of neighbouring sarcomeres and provides structural integrity. α-Actinin-2 also binds to titin, which is connected to thick filaments. This interaction is regulated by a phosphoinositide-based mechanism [7, 8]. However, many other details of the molecular architecture of the Z-disc remain elusive, and many questions still need to be addressed. For example, why is tropomyosin distributed all along thin filaments of the sarcomere with exception of the Z-disc [6]? How are interactions of α-actinin with numerous binding partners translated into Z-disc assembly?

The Z-disc component myotilin is involved in multiple interactions by directly binding to α-actinin-2, filamin C, FATZ/myozenin/calsarcin, ZASP/cypher and F-actin [9–13]. It was therefore identified as a key structural component of Z-discs and proposed to control sarcomere assembly [9–11]. Mutations in the human myotilin gene are associated with myofibrillar myopathy (MFM) [14, 15], supporting the notion that myotilin is important for proper maintenance, organization and/or function of the Z-disc.

Myotilin is a member of the palladin/myopalladin/myotilin family and contains two Ig domains that are involved in myotilin dimerization, and interact with F-actin and filamin C (**Figure 1B**) [9, 10, 16, 17]. The shortest fragment of myotilin that can bind F-actin includes the two Ig domains preceded by a stretch of intrinsically disordered residues (aa 214-442) [17]. A longer myotilin fragment including N- and C-terminal regions flanking two Ig domains (aa 185-498) can cross-link actin filaments *in vitro* and *in vivo* [17]. In addition, full-length protein prevents F-actin disassembly induced by latrunculin A [10], revealing its role in stabilization and anchoring of thin filaments in the Z-discs [10]. Myotilin dimerization was proposed to be important for F-actin cross-linking [10]. The shortest myotilin fragment with the ability to dimerize includes the Ig2 domain and the C-terminal tail (aa 345-498) [17]. Although an anti-parallel myotilin dimer was suggested [10] the insight into molecular determinants of myotilin dimerization remains limited.

To elucidate the molecular mechanism of myotilin interaction with F-actin and the role of this interaction in sarcomeric Z-disc assembly and structure, we used an integrative structural biology approach to build the first structural model of the myotilin:F-actin complex, which could serve as a blueprint for interaction of Ig-domain containing proteins with F-actin *via* two or more consecutive domains, such as the entire palladin family and filamins [18–21]. This structural model suggests that binding sites for myotilin, tropomyosin and α-actinin-2 on F-actin partially overlap, which we confirmed by competition assays. Furthermore, we characterized the interaction of myotilin with α- actinin-2 and showed that PI(4,5)P_2_ does not have a direct regulatory effect on myotilin.

Based on our results we propose a model in which myotilin simultaneously binds F-actin and α- actinin with the concomitant displacement of tropomyosin. This renders myotilin not only structural support for Z-disc architecture but also a co-organiser of Z-disc assembly. Furthermore, our structural model of full-length myotilin provides a platform for understanding the molecular basis of disease-causing mutations and their impact on the structure, ligand binding and/or regulation.

## Results

### Myotilin is a conformational ensemble in solution

The previously performed disorder tendency analysis of full-length myotilin revealed that the N-terminal region of myotilin displays characteristics of an intrinsically disordered protein (IDP) [22]. The meta-structure analysis, which reflects how likely a specific residue is to be located within a compact 3D structure, was carried out as described previously [23], and indicated a tendency for local compactions in the N-terminal ID region (**Figure 1B**).

For characterization of full-length myotilin, we fused thioredoxin (Trx) to its N-terminus (Trx-MYOT) to prevent progressive N-terminal degradation. A globular protein at the N-terminus of myotilin was shown not disturb its function *in vivo*, where GST, Myc and GFP fusions have been extensively used in cell biophysics experiments [13, 24, 25]. To investigate the hydrodynamic and structural properties of Trx-MYOT, we first performed size exclusion chromatography coupled to SAXS (SEC-SAXS) (**Figure S1A**). Right-angle laser light scattering (RALLS) data collected in parallel confirmed that the elution peak corresponded to a monomeric protein (**Figure S1A**). The overall molecular parameters (molecular weight M_W_, the radius of gyration, R_g_, and maximal intermolecular distance, D_max_,) derived from the resulting SAXS profile are also compatible with a monomeric species (**Figure 1C, Table S1**). The featureless descent of the SAXS profile in log plot (**Figure 1C**), and the plateau in the dimensionless Kratky plot (**Figure S1B**) are characteristic for the scattering of particles that are disordered and/or flexible. The calculated atom-pair distance distribution functions *P*(*r*) skewed to shorter distances and the D_max_ value of ∼ 200 Å also suggest flexible and elongated “string like” structures (**Figure 1C, inset**).

The derived *ab initio* molecular envelope of Trx-MYOT displayed an elongated multi-subunit shape (**Figure 1D**). To better account for the flexibility of Trx-MYOT, we employed the ensemble optimization method (EOM) [26]. A genetic algorithm-based selection of representative models with their respective volume fractions in solution resulted in the best fit to the data with χ^2^=1.01 (**Figures 1C and 1E**). Overall, the EOM-selected models tended to be more compact than those in the random pool. This is visible from the shift to smaller values of D_max_ and R_g_ derived from the selected models compared to the distribution derived from the original pool, as the most extended models were not selected (**Figures 1E, S1C and S1D**). Based on this analysis, Trx-MYOT displays a tendency to occupy more compact confirmations compared to a chain with random coil linkers, in line with the predicted compactness (**Figure 1B**) and the disorder tendency analysis [22].

In summary, Trx-MYOT in solution displays a flexible conformational ensemble with local clusters of compactness in the intrinsically disordered regions (IDR), in addition to the structured Ig domains connected by flexible linkers.

### Myotilin forms concentration-dependent dimers

Although myotilin was suggested to form antiparallel dimers [10] full-length myotilin was monomeric under our experimental conditions. The shortest myotilin fragment with the ability to dimerize was reported to include the Ig2 domain and the C-terminal tail [17]. To investigate molecular determinants of dimerization we used SAXS on a series of truncated constructs — Ig1Ig2^250-444^ (Ig1Ig2 tandem), Ig1Ig2^220-452^ and Ig1Ig2^250-498^— due to the inherently low solubility of Trx-MYOT.

We monitored the maximum scattering intensities extrapolated to zero angle (I_0_) for a concentration series of the Ig1Ig2^220-452^ construct, and observed a concentration-dependent increase in the average molecular mass of resulting particles (**Figure 1F, inset**). Accordingly, the *P*(*r*) function for Ig1Ig2^220-452^ showed a transition from a typical distribution for an extended two-domain protein, with a significant peak at approximately 25 Å and a minor at 50 Å, to a distribution with increased frequencies of the 50 Å peak and a new minor signal at in the region 80-100 Å (**Figure 1F**). Here, the shortest vector corresponds to the distances within Ig domains, while the second and the third peaks correspond to inter-domain distances within and between the subunits with concomitant increase of D_max_ value to approximately 150 Å (**Figure 1F, and Table S1**). The notable increase of the D_max_ value in a concentration dependent manner from 110 Å to 150 Å together with aforementioned features of the *P(r)* suggests an antiparallel staggered dimer formed either by interactions *via* Ig1 (head-to-head) or *via* Ig2 domains (tail-to-tail), or a staggered parallel dimer *via* Ig1-Ig2. The calculated structural parameters (D_max_ and *P(r)*) of the non-staggered parallel or antiparallel dimer are not in line with the experimentally derived values, thus making such architecture less likely (**Figures 1F, S1E and S1F**).

*P(r)* functions of Ig1Ig2^250-444^, Ig1Ig2^220-452^ and Ig1Ig2^250-498^ derived from scattering data at comparable concentrations showed that Ig1Ig2^220-452^ and Ig1Ig2^250-498^ displayed a high second peak at 50 Å, and corresponding D_max_ values of approximately 150 Å (**Figure 1G, and Table S1**), whereas Ig1Ig2^250፧444^ did not. Guinier region analysis revealed an apparent increase of R_g_ for the Ig1Ig2^220-452^ and Ig1Ig2^250-498^, whereas Ig1Ig2^250፧444^ displayed molecular parameters of a monomeric species, comparable to those observed for Ig1Ig2^220፧452^ at low concentrations (**Figures 1F, and Table S1**). Consequently, M_W_ values calculated from I_0_ using the Guinier approximation correspond to the expected M_W_ of a dimer for Ig1Ig2^220-452^, approaching dimer for Ig1Ig2^250-498^ and of a monomer for Ig1Ig2^250-444^ (**Table S1**), corroborating the role of regions flanking the Ig1Ig2.

R405 is mutated in muscular dystrophy, and was suggested to be responsible for defective homo-dimerization of myotilin [27]. We therefore assessed the dimerisation propensity of Ig1Ig2^220-452 R405K^ mutant using SAXS. Comparative analysis of SAXS data and derived molecular parameters (D_max_, R_g_, *P(r)*, M_W_) revealed that the conserved R405K mutation which preserves the positive charge does not impair dimer formation *in vitro*, (**Figures S1G and Table S1**), in contrast to previously reported defective homo-dimerisation observed by immunoprecipitation and yeast two hybrid screens [27].

Our results together with published data on myotilin dimerization collectively show that: (i) N- and C-terminal regions flanking the Ig1Ig2 domains contribute to dimer stabilisation in solution; (ii) the fully dimeric form is observed only at high protein concentrations (> 1.0 mM), suggesting a weak association constant, explaining why only monomers of Trx-MYOT were observed in our SEC-SAXS experiments, where only low concentrations of Trx-MYOT could be used due to its aforementioned low solubility; (iii) tail-to-tail dimerisation reconciles both SAXS analysis and proposed mode of incorporation in the Z-disc, ochestered by interactions with filamin C and α-actinin-2 [10].

### Tandem Ig-domains of myotilin together with contiguous regions are required for high affinity binding to F-actin

To characterise myotilin:F-actin interaction and to elucidate the suggested role of segments flanking Ig1Ig2 [10, 17], we determined the binding affinity of full-length Trx-MYOT, and a series of truncated constructs, using F-actin co-sedimentation assays (**Figure 2A**).

**Figure 2.**
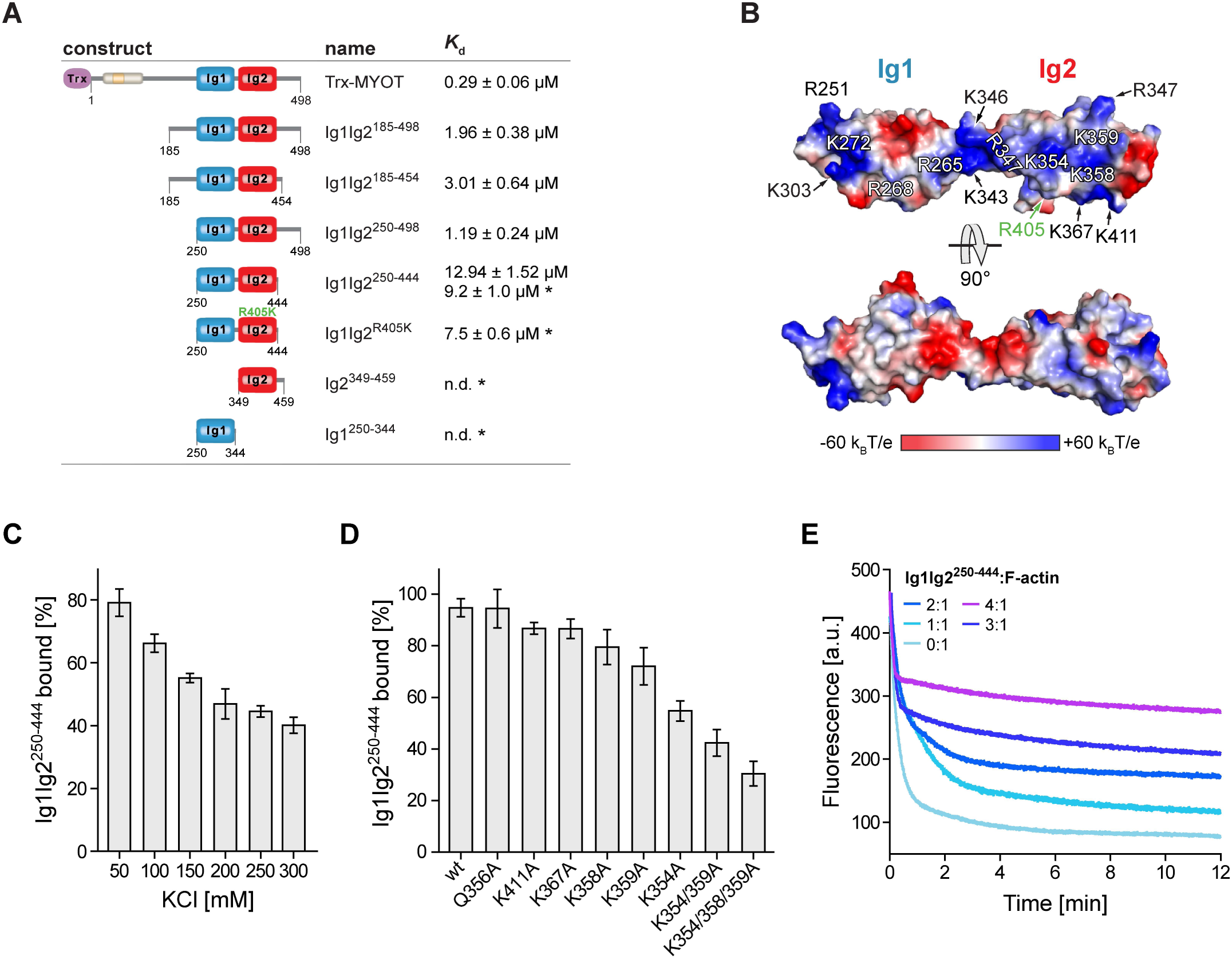
Myotilin influences F-actin dynamics by binding *via* its Ig domains and disordered flanking regions. (**A**) Summarized results of the actin co-sedimentation assays. The asterisk denotes values obtained in assays performed under B1 conditions (for details see Material and Methods). (**B**) Surface electrostatic potential mapped on the Ig1Ig2 structural model [22]. Surfaces are coloured on a red-white-blue gradient, as calculated by Adaptive Poisson-Boltzmann Solver [83]. Residues forming a basic patch (blue) on both Ig domains are labelled. The residue (R405) involved in muscular dystrophy-causing mutation R405K is shown in green [27]. (**C**) Binding of Ig1Ig2^250-444^ to F-actin at increasing salt conditions. Graph data are presented as a percentage of theoretical total binding (100 %) corresponding to the total amount of protein (8 µM) used in each experiment. Data represent mean values ± SEM of three independent experiments. The mean binding of Ig1Ig2^250-444^ to F-actin in 50 mM KCl was significantly different to the means for all other tested concentrations of KCl. Significance was assessed using one-way ANOVA with Holm-Sidak test, p < 0.001 for salt concentration of 50 mM vs. all other tested concentrations. (**D**) Binding of Ig1Ig2^250-444^ and its mutant versions to F-actin. Graph data are presented as in (**C**). Data represent mean values ± SEM of three independent experiments. The mean binding to F-actin of all mutants except Q356A, K411A and K367A were significantly different from the means of WT Ig1Ig2^250-444^. Significance was assessed using one-way ANOVA with Holm-Sidak test, p < 0.01 for WT vs. K358A and p < 0.001 for WT vs. all other mutants. (**E**) Effects of Ig1Ig2^250-444^ on F-actin depolymerization. Actin filaments were depolymerized in the presence or absence of Ig1Ig2^250-444^ at molar ratios indicated in the figure. See also **Figure S2**.

In the initial experiments (using conditions B1, for details, see Materials and Methods) with single Ig domains (Ig1^250-344^ and Ig2^349-459^), and their tandem (Ig1Ig2^250-444^), neither Ig1, nor Ig2 alone displayed significant binding to F-actin, while Ig1Ig2^250-444^ bound to F-actin with *K*_d_ = 9.2 ± 1.0 µM (**Figures 2A and S2A**). When the same experimental conditions were used for Trx-MYOT, its self-pelleting was observed making interpretation of the data difficult. To reduce the fraction of self-pelleted Trx-MYOT, we optimized the conditions for the co-sedimentation assays using a differential scanning fluorimetry (DSF)-based pH-screen (**Figure S2C**). We found that a slightly acidic pH, i.e. an adjustment of the assay buffer from pH 7.4 (in conditions B1) to pH 6.8 (conditions B2, for details, see Materials and Methods) rendered Trx-MYOT more stable as well as more soluble. To assess the effect of pH (6.8), affinity of Ig1Ig2^250-444^ to F-actin was determined using the conditions B2, too. The observed apparent *K*_*d*_ (12.9 ± 1.5 µM) was in good agreement with the affinity measured in the B1 conditions (*K*_*d*_ = 9.2 ± 1.0 µM) (**Figures 2A and S2B**, left panel), suggesting that diverse conditions used do not have a major effect on the myotilin affinity to F-actin.

The apparent *K*_*d*_ of Trx-MYOT for F-actin in conditions B2 is 0.29 ± 0.06 µM (**Figures 2A and S2B**, right panel), indicating relatively strong binding compared to other F-actin binding proteins of the palladin family. For instance, the affinity of full-length palladin to F-actin was reported to be 2.1 ± 0.5 µM [18]. To further delineate the role of regions flanking Ig1Ig2, Ig1Ig2^185-454^, Ig1Ig2^250-498^, and Ig1Ig2^185-498^ were assayed for F-actin binding (**Figures 2A and S2B**). All constructs bound to F-actin with similar affinity (*K*_*d*_ = 3.0 ± 0.6 µM, 1.2 ± 0.2 µM, and 2.0 ± 0.4 µM for Ig1Ig2^185-454^, Ig1Ig2^250-498^, and Ig1Ig2^185-498^, respectively), and thus significantly stronger than the Ig1Ig2^250-444^ encompassing only the two Ig domains (**Figures 2A and S2B**).

The surface electrostatic potential of the tandem Ig1Ig2 domains displays positive clusters on one face of the domains (‘BED’ β-sheet) and in general a more pronounced basic character of Ig2 in contrast to Ig1 (**Figure 2B**). Positive clusters have also been observed in several other F-actin binding proteins, including palladin [28], suggesting an electrostatically driven interaction. To investigate the influence of ionic strength on the affinity we performed co-sedimentation assays with increasing NaCl concentrations. As shown in **Figure 2C**, higher salt concentrations reduced the ability of myotilin to co-sediment with F-actin.

To further corroborate the electrostatic nature of the interaction, we gradually neutralized the positive charge by mutating the basic residues of the Ig2 domain to Ala in the construct Ig1Ig2^250-444^. While mutations of Q356A, K411A and K367A did not significantly affect binding to F-actin compared to the wild-type, single mutations K358A, K359A, and K354A had a negative effect, which was remarkably potentiated in the double K354/359A mutant, and even more in the K354/358/359A triple mutant (**Figure 2D**). The latter mutant showed a 70% reduction in binding to F-actin, suggesting that this basic patch in Ig2 plays an important role in F-actin binding (**Figure 2B and 2D**).

The muscular dystrophy-causing mutant R405K (Ig1Ig2^R405K^) [27] was also tested, but did not show any notable difference in F-actin affinity with respect to the wild-type (**Figures 2A and S2A**), which agrees with its lateral location with respect to the identified positively charged patch involved in F-actin binding. Furthermore, the mutation retains the positive charge at this position (**Figure 2B**).

Since binding data is available for other actin-binding Ig domains, e.g. for Ig domains of palladin and filamin A [19], we conducted a comparative structural analysis of the actin-binding Ig domains of myotilin, the Ig3 domain of palladin, and Ig10 of filamin A, which revealed charged residues positioned on the same face in all analysed Ig domains (**Figures S2D**). Notably, these residues coincide with those indicated by co-evolution analysis [29] as potentially functional sites (**Figure S2D**, lower panel). Specifically, lysine residues K1008, K1011 and K1044 on palladin Ig3 domain were shown to be essential for F-actin binding [28]. Equivalent residues are also present in myotilin (**Figure S2D**, upper panel), and form a basic patch, which extends from one Ig domain to the other (**Figure 2B**).

Finally, we biochemically characterized the previously observed myotilin binding to (monomeric) G-actin [17] using a constitutively monomeric mutant of actin, DVD-actin [30]. We employed microscale thermophoresis (MST) with various myotilin tandem or isolated Ig domain constructs. The affinity of myotilin constructs to non-polymerizable DVD-actin was two orders of magnitude lower than that to F-actin (**Figures S2E and S2F**). The highest affinity displayed by construct Ig1Ig2^185-454^ indicates that, similar to F-actin, the Ig-domain-flanking regions could be involved in modulating actin dynamics.

We hence performed depolymerization assays, to obtain further insight into the role of myotilin in actin dynamics. As also observed for the full-length myotilin [10], addition of Ig1Ig2^250-444^ reduced F-actin depolymerization rate in a concentration-dependent manner (**Figure 2E**), suggesting that the tandem Ig domains play the critical role in actin binding.

In summary, our results reveal that myotilin has a role in stabilization of actin filaments and actin dynamics, where the tandem Ig domains play the central role and that both the N- and C-terminal neighbouring regions further enhance affinity and stabilise the myotilin:F-actin interaction (**Figure 2A**).

### Integrative structural model of the myotilin:F-actin complex

To validate and identify potential new contacts with Ig domains and with their contiguous regions, as implied from the binding and mutational data (**Figure 2A and 2D**), we performed XL-MS experiments on Ig1Ig2^250-444^, Ig1Ig2^185-454^ and Ig1Ig2^185-498^ bound to F-actin (**Figures S3A-S3C**). As expected, specific cross-links involving both Ig1 and Ig2 domains as well as their N- and C-terminal flanking regions (**Figures 3A and Table S2**) were found. While only one cross-link was detected between the Ig1 domain and F-actin (K303 on Ig1 and D27 on actin), there were several cross-links between the Ig2 domain and F-actin. This is in line with its higher affinity and the charged nature of this interaction (**Figures 2B, 2D, and S2A**), leading to cross-links of basic lysine residues on Ig2 domain with the acidic residues located in subdomain 1 of actin. Of the Ig2 domain residues cross-linked to F-actin, we also found K354 involved in the interaction using mutagenesis and co-sedimentation assays (**Figures 2D and 3A**). Furthermore, the cross-links between acidic myotilin residues (D236, D239 and D241) and basic K328 and K330 on actin (**Figure 3A and Table S2**) map to the region preceding myotilin Ig1 in constructs Ig1Ig2^185-454^ and Ig1Ig2^185-498^. The latter construct yielded additional cross-links between positively charged K452, K462, K469 and K474 mapping to the region C-terminal to Ig2, and negatively charged residues E336, D27 and D26 on actin (**Figure 3A and Table S2**).

**Figure 3.**
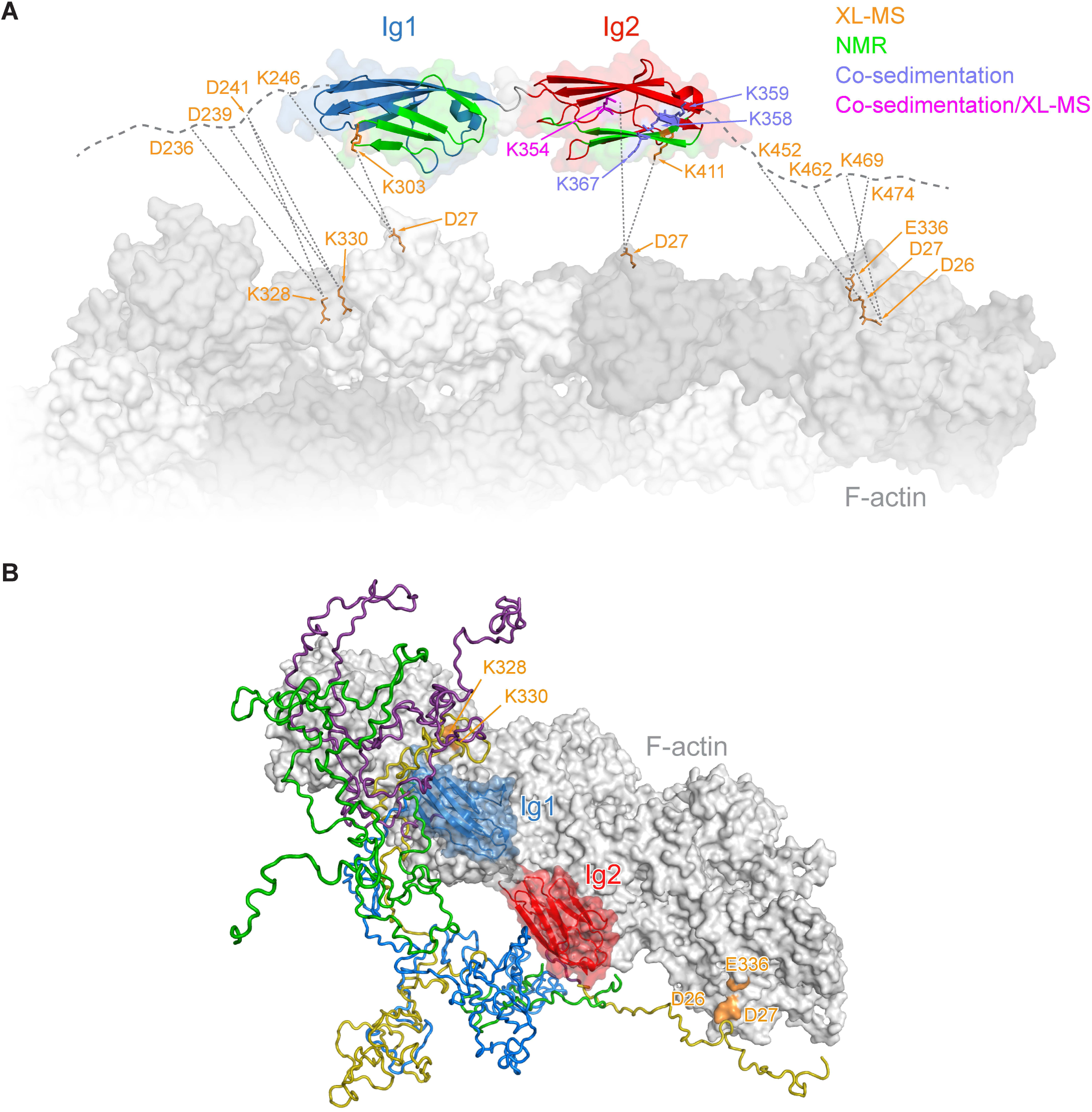
Integrative structural model of myotilin:F-actin complex. (**A**) Summary of experimental data on myotilin:F-actin complex, obtained from XL-MS (orange), co-sedimentation/mutagenesis analysis (medium slate blue), and NMR (green). K354 was found to be involved in the interaction by both XL-MS and co-sedimentation/mutagenesis analysis (magenta). Thick, grey dashed lines represent N- and C-terminal flanking regions of myotilin Ig domains. Selected residues found in cross-links between actin and myotilin are shown in orange. (**B**) Model of the full-length myotilin bound to F-actin. Ig domains of myotilin were docked on F-actin and combined with flexible (not docked) models of N- and C-terminal parts of the protein from EOM analysis of Trx-MYOT (**Figure 1E**). The four most populated conformations of N- and C-terminal parts are shown in distinct colours. Selected residues of actin found in cross-links with myotilin are shown in orange. See also **Figure S3 and Table S2**.

To construct an integrative molecular model of the myotilin:F-actin complex, we combined our structural model of Trx-MYOT (**Figure 1E**) and experimental data derived from *in vitro* mutational analysis and XL-MS, which we used as experimental distance restraints that guided the macromolecular docking using Haddock 2.2 [31] (for details see Material and Methods). In this model the tandem Ig domains land on the subdomains 1 of the adjacent actin subunits, with the flexible N- and C-terminal regions extending to the binding sites on neighbouring actin subunits as suggested by XL-MS data (**Figures 3B**), supporting and explaining published data [10, 17].

NMR was subsequently used to validate the structural model of myotilin:F-actin complex. NMR spectra of the isolated ^15^N-labelled Ig1 and Ig2 domains showed similar characteristics to those available in the BMRB databank (Ig1: 7113; Ig2: 16370) [32], and hence *de novo* assignment was not required in order to obtain a subset of assigned peaks. Construct Ig1Ig2^250-444^ also showed a similar peak pattern, allowing the use of assignments of the single domains to obtain a partial assignment of this tandem construct. In the initial HSQC experiments, ^15^N-labelled Ig1^250-344^ was titrated with F-actin, which induced a reduction of the signal intensities (**Figure S3D**). Measurements of the ^15^N-labelled Ig2 domain (Ig2^349-459^) showed a stronger reduction in signal intensity upon addition of F-actin compared to those carried out with the Ig1 domain (**Figure S3D**) indicating stronger binding of Ig2 to F-actin.

Further experiments with ^15^N-labelled Ig1Ig2^250-444^ showed a shift pattern upon addition of F-actin, where the regions with significant HSQC shifts are: 257-270, 296-309, 363-374 and 405-413 (**Figure S3E and S3F**) with segment 301-306 unfortunately not being represented due to low signal intensity [32]. These regions map to the ‘BED’ β-sheet of both Ig domains and additionally to the β-strand A’ in Ig1, and are in agreement with the interaction sites mapped by binding, mutational and XL-MS data (**Figure 2A, 2D and 3A**).

### Myotilin competes with tropomyosin and α-actinin-2 for the binding sites on F-actin

The actin residues identified in the myotilin:F-actin interaction by XL-MS include K328, K330 and D26, D27 and E336 (**Figures 3A and 3B**). Interestingly, K328, together with neighbouring K326, form the major interaction site for tropomyosin on F-actin [33], implying that myotilin and tropomyosin could compete for binding to F-actin. We therefore created a structural model of myotilin bound to F-actin in the presence of tropomyosin (**Figure 4A**). In this model, two Ig domains of myotilin only marginally overlap with tropomyosin bound to F-actin. However, their N- and C-terminal flanking regions found by XL-MS analysis to interact with actin residues (D26, D27, K326, K328, and E336), map to or close to the tropomyosin binding site (**Figure 4A**), suggesting that they might interfere with F-actin:tropomyosin binding.

**Figure 4.**
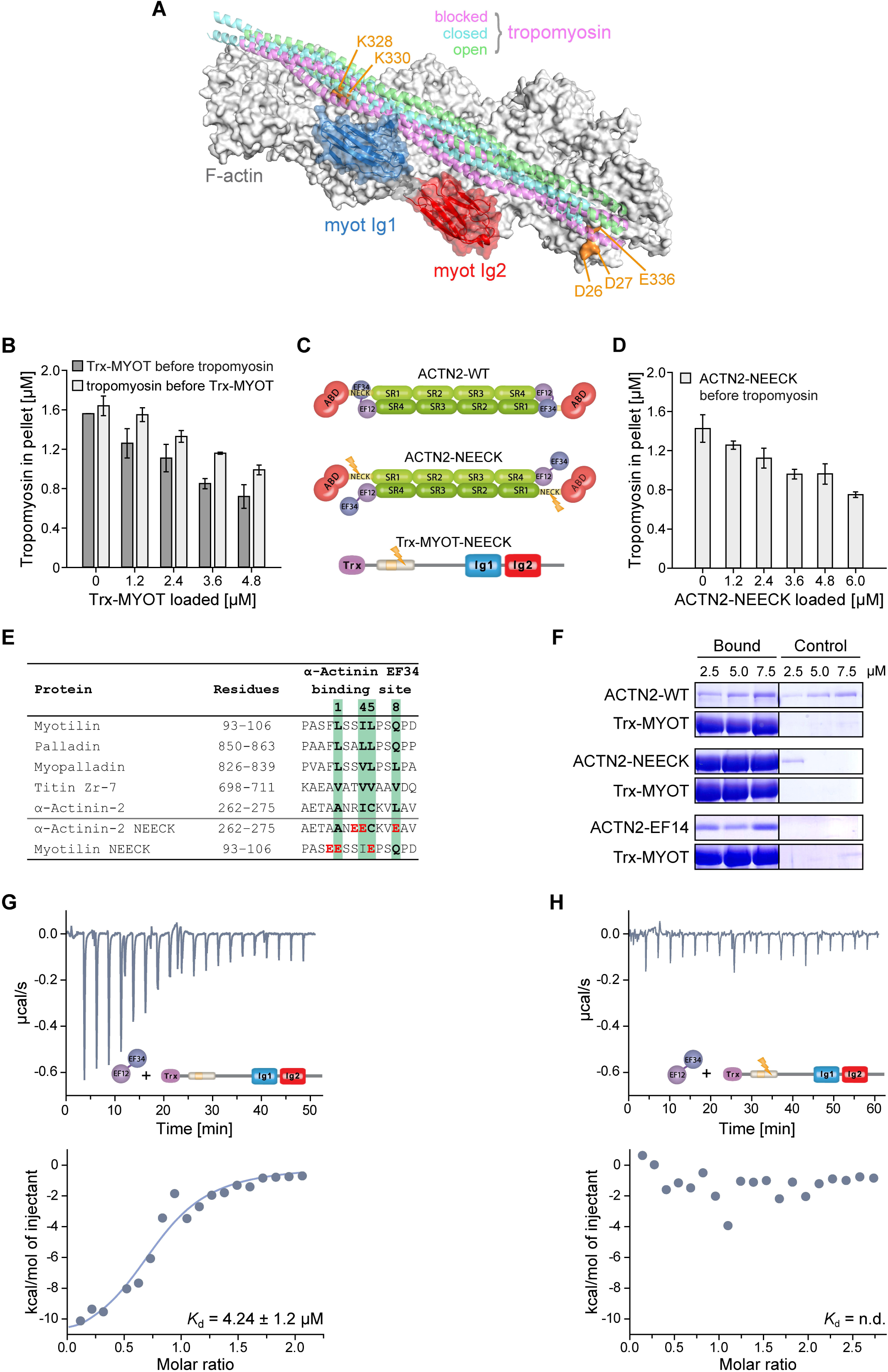
Myotilin regulates binding of tropomyosin to F-actin and interacts with α-actinin-2. (**A**) Model of tandem Ig domains of myotilin bound to F-actin superimposed with the cryo-EM structures of cardiac tropomyosin bound to F-actin in the blocked (magenta ribbons, PDB: 5NOG), closed (cyan ribbons, PDB: 5NOL), and open (green ribbons, PDB: 5NOJ) structural states [84]. Residues of actin found in cross፧links with N- and C-terminal regions flanking myotilin Ig domains are shown in orange (see **Figure 3A**). (**B**) Effects of myotilin on tropomyosin:F-actin interaction. Myotilin (Trx-MYOT) at increasing concentrations was added to F-actin before (Trx-MYOT before tropomyosin) or after (tropomyosin before Trx-MYOT) incubation of F-actin with the fixed amount of tropomyosin. Mean values (± SEM) of three independent experiments are shown. For “Trx-MYOT before tropomyosin” the mean binding of tropomyosin to F-actin at all concentrations of Trx-MYOT was significantly different from the means in the absence of Trx-MYOT. Significance was assessed using one-way ANOVA with Holm-Sidak test, p < 0.01 for 0 vs. 1.2 μM Trx-MYOT and p < 0.001 for 0 vs. all other concentrations of Trx-MYOT. For “Tropomyosin before Trx-MYOT” the mean binding of tropomyosin to F-actin at all concentrations of Trx-MYOT, except for 1.2 μM Trx-MYOT, was significantly different from the means in the absence of Trx-MYOT. Significance was assessed using one-way ANOVA with Holm-Sidak test, p < 0.05 for 0 vs. 2.4 μM Trx-MYOT, p < 0.01 for 0 vs. 3.6 μM Trx-MYOT and p < 0.001 for 0 vs. 4.8 μM Trx-MYOT. (**C**) Schematic presentation of the wild-type α-actinin-2 (ACTN2-WT), its constitutively open mutant (ACTN2-NEECK), and mutant of Trx-MYOT (Trx-MYOT-NEECK), possessing mutations resembling those in the ACTN2-NEECK (see **Figure 4E**). Lightning bolt depicts position of mutations. (**D**) Effects of myotilin on α-actinin:F-actin interaction. ACTN2-NEECK was added to F-actin at increasing concentrations before (ACTN2-NEECK before tropomyosin) incubation of F-actin with the fixed amount of tropomyosin. Mean values (± SEM) of three independent experiments are shown. The mean binding of tropomyosin to F-actin at all concentrations of ACTN2-NEECK was significantly different from the means in its absence. Significance was assessed using one-way ANOVA with Holm-Sidak test, p < 0.05 for 0 vs. 1.2 μM ACTN2-NEECK, p < 0.01 0 vs. 2.4 μM ACTN2-NEECK, and p < 0.001 for 0 vs. all other concentrations of ACTN2-NEECK. (**E**) Sequence alignment of proteins and their residues involved in binding to α-actinin EF34. Residues of the CaM-binding motif 1-4-5-8 are shown in bold and boxed in green. Residues mutated in ፧-actinin-2 NEECK and myotilin NEECK are shown in red. (**F**) Binding of myotilin to α-actinin. ACTN2-WT, ACTN2-NEECK, for details see (**C** and **E**), and α-actinin-2 EF14 (ACTN2-EF14) were subjected at increasing concentrations to pull-down assay with (bound) or without (control) Trx-MYOT. (**G** and **H**) Results of ITC experiments quantifying the interaction between (**G**), Trx-MYOT, or (**H**) its mutant (Trx-MYOT-NEECK), for details see (**C** and **E**) and ACTN2-EF14. n.d., not determined. See also **Figure S4**.

To test this hypothesis, we performed competition co-sedimentation assays where F-actin was incubated with increasing concentrations of Trx-MYOT, either before or after adding a fixed amount of the striated muscle isoform of human tropomyosin (Tpm1.1). The amount of tropomyosin bound to F-actin decreased with increasing concentrations of Trx-MYOT (**Figures 4B and S4A**), uncovering that myotilin competes with tropomyosin for F-actin binding and displaces tropomyosin from F-actin.

In the myofibrillar Z-disc, myotilin associates with the striated muscle-specific isoform α-actinin-2 [9, 11], the major Z-disc protein. As the stress-fibre-associated tropomyosin isoforms were shown to compete with non-muscle α-actinin-1 and α-actinin-4 for F-actin binding [34, 35], we examined whether α-actinin-2 is also able to compete with tropomyosin for F-actin. We used the constitutively open variant of α-actinin-2, in which mutations abrogating binding of EF34 to the α-actinin-2 “neck” region were introduced (ACTN2-NEECK, **Figure 4C**) to resemble the Z-disc bound conformation of α-actinin-2 [7]. We performed competition co-sedimentation assays, in which α-actinin-2 was incubated at increasing concentrations with F-actin before the addition of tropomyosin. As expected, binding of α-actinin-2 to F-actin prevented tropomyosin-F-actin binding in a concentration dependent manner (**Figures 4D and S4B**).

Comparative structural analysis of the myotilin:F-actin:tropomyosin model (**Figure 4A**) and the cryo-EM structures of F-actin decorated with the actin binding domain (ABD) of α-actinin-2, spectrin or filamin [36–38], showed that binding sites of myotilin, tropomyosin and α-actinin-2 on F-actin overlap (**Figure S4C**). Furthermore, comparison of binding affinities of myotilin, tropomyosin and α-actinin-2 to F-actin [34, 39–42] together with our competition assays (**Figures 4B and 4D**), indicate that myotilin could displace both tropomyosin and α-actinin-2 from F-actin, suggesting a novel role of myotilin as a regulator of access of tropomyosin and other actin-binding proteins to F-actin in the Z-disc.

### Myotilin binds α-actinin-2 using the same pseudoligand regulatory mechanism as titin

α-Actinin-2 interacts with titin as well as with pallading family of proteins *via* its C-terminal EF-hands (EF34) of the calmodulin-like domain (CAMD) [43–45]. The interaction between α-actinin-2 and titin binding motifs, the Z-repeats, has been well characterised [7, 8, 43], and was shown to be regulated by an intramolecular pseudoligand mechanism, in which a titin Z-repeat-like sequence (“neck”, which contains the 1-4-5-8 binding motif) connecting the ABD and the first spectrin repeat of α-actinin-2, interacts with EF34 of the juxtaposed CAMD (**Figure 4C**). In this closed conformation, EF34 are not available for interactions to titin Z-repeats unless PI(4,5)P_2_ catalyzes conformational switch to the open confirmation through their release from the “neck”, and hence activating titin binding [7].

The interaction between α-actinin-2 and myotilin maps to the N-terminal part of myotilin, which also contains the conserved 1-4-5-8 motif (residues 95-106, **Figures 4C and 4E**) similar to palladin and myopalladin [45]. This was suggested by NMR experiments where a myotilin peptide containing the binding motif (**Figure 4E**) was titrated to EF34 [45].

To characterize the interaction of myotilin with α-actinin-2 with full-length proteins we firstly performed pull-down assays using either wild-type α-actinin-2 (ACTN2-WT), its constitutively open mutant ACTN2-NEECK, or CAMD encoding all four EF-hands of α-actinin-2 (ACTN2-EF14), with Trx-MYOT (**Figures 4C and 4E**). We observed myotilin binding to ACTN2-NEECK and ACTN2-EF14, but not to ACTN2-WT (**Figure 4F**) indicating that opening of α-actinin-2 is necessary for its interaction with myotilin, as was also revealed for titin [7].

To validate the binding site between α-actinin-2 and myotilin (**Figures 4C and 4E**), we next performed ITC experiments with ACTN2-EF14, Trx-MYOT and its mutant variant (Trx-MYOT-NEECK), where mutations resembling those in ACTN2-NEECK were introduced in the binding motif to disrupt the interaction. Indeed, the affinity of Trx-MYOT to α-actinin-2 was found to be in µM range (*K*_*d*_ = 4.2 ± 1.2 μM, **Figure 4G**), whereas no binding of Trx-MYOT-NEECK to ACTN2-EF14 was observed (**Figure 4H**).

Altogether our data reveal that myotilin, which contains the conserved α-actinin-2 binding motif, binds to the open confirmation of α-actinin-2 *via* pseudoligand regulatory mechanism.

### Interaction of myotilin with F-actin is not regulated by PI(4,5)P_2_

The Ig3 domain of palladin is the minimum fragment necessary for binding to F-actin [18, 28]. Furthermore, Ig3 interacts with the head group of PI(4,5)P_2_ with a moderate affinity (*K*_*d*_ = 17 μM) leading to a decreased F-actin cross-linking and polymerization activity [46]. Myotilin binding to α-actinin-2 follows the same PI(4,5)P_2_ mechanism as titin, while its F-actin binding is similar to that of palladin. Therefore, we investigated whether myotilin binds PI(4,5)P_2_ as well, using a liposome co-sedimentation assay [46]. The amount of protein bound to liposomes was examined by using a constant concentration of Ig1Ig2^250-444^, while varying the PI(4,5)P_2_ concentration (0-20%) in 1-palmitoyl-2-oleoyl-sn-glycero-3-phosphocholine (POPC) vesicles. As positive controls, double C2-like domain-containing protein beta (Doc2b), a known sensor of PI(4,5)P_2_ [47], and the Ig3 domain of palladin were used (**Figures S4D and S4E**). Both proteins, but not myotilin Ig1Ig2^250-444^, co-pelleted with PI(4,5)P_2_-POPC vesicles in a PI(4,5)P_2_ concentration dependent manner (**Figures S4D and S4E**) indicating that myotilin does not interact with PI(4,5)P_2_. This observation was further corroborated by structure-based sequence analysis, which showed that residues important for binding of PI(4,5)P_2_ to palladin [46] are not present in the Ig2 domain of myotilin (**Figure S4F**). Therefore, the myotilin interaction with F-actin is most likely not regulated by PI(4,5)P_2_.

### Binding to F-actin affects myotilin mobility and dynamics in cells

To assess whether actin-binding affects the cellular dynamics of myotilin, we expressed N-terminal EGFP fusion of myotilin, or its mutants that showed impaired F-actin binding activity *in vitro* (K354A, K359A, K354/359A, K354/358/359A), in differentiating C2C12 mouse myotubes. After seven days of differentiation, all of the tested mutants and the wild-type myotilin showed a similar distribution and localized to Z-discs (**Figure 5A, panel “pre-bleaching”**), suggesting that similarly to EGFP, N-terminal Trx-tag used for *in vitro* studies does not perturb myotilin function. Fluorescence recovery after photobleaching (FRAP) was performed to compare the mobility and dynamics of the mutant variants with wild-type myotilin (**Figure 5A**). Wild-type myotilin showed rapid dynamics with 80% fluorescence recovery after 300 s (median halftime t_1/2_ of 72.4 s) (**Figure 5B**), which is in excellent agreement with previously published reports [48, 49]. While the single mutant K359A only showed slight, non-significant changes (t_1/2_ 64.1 s), K354A showed significantly faster recovery with t_1/2_ of 48.2 s (**Figures 5B and 5C**). Protein dynamics further increased proportionally to the number of mutations in double (t_1/2_ 35.5 s) and triple (t_1/2_ 32.3 s) mutants (**Figures 5B and 5C**). The differences in recovery times are evident by comparing the fluorescence signals in **Figure 5A**: the wild-type and K359A myotilin hardly showed any recovery after 5 or even 20 s, indicating relatively slow dissociation and high affinity, but a clear signal can be seen for the other three mutants with reduced actin binding strength. At the same time, the mobile fraction of the mutants increased correspondingly (**Figure 5D**).

**Figure 5.**
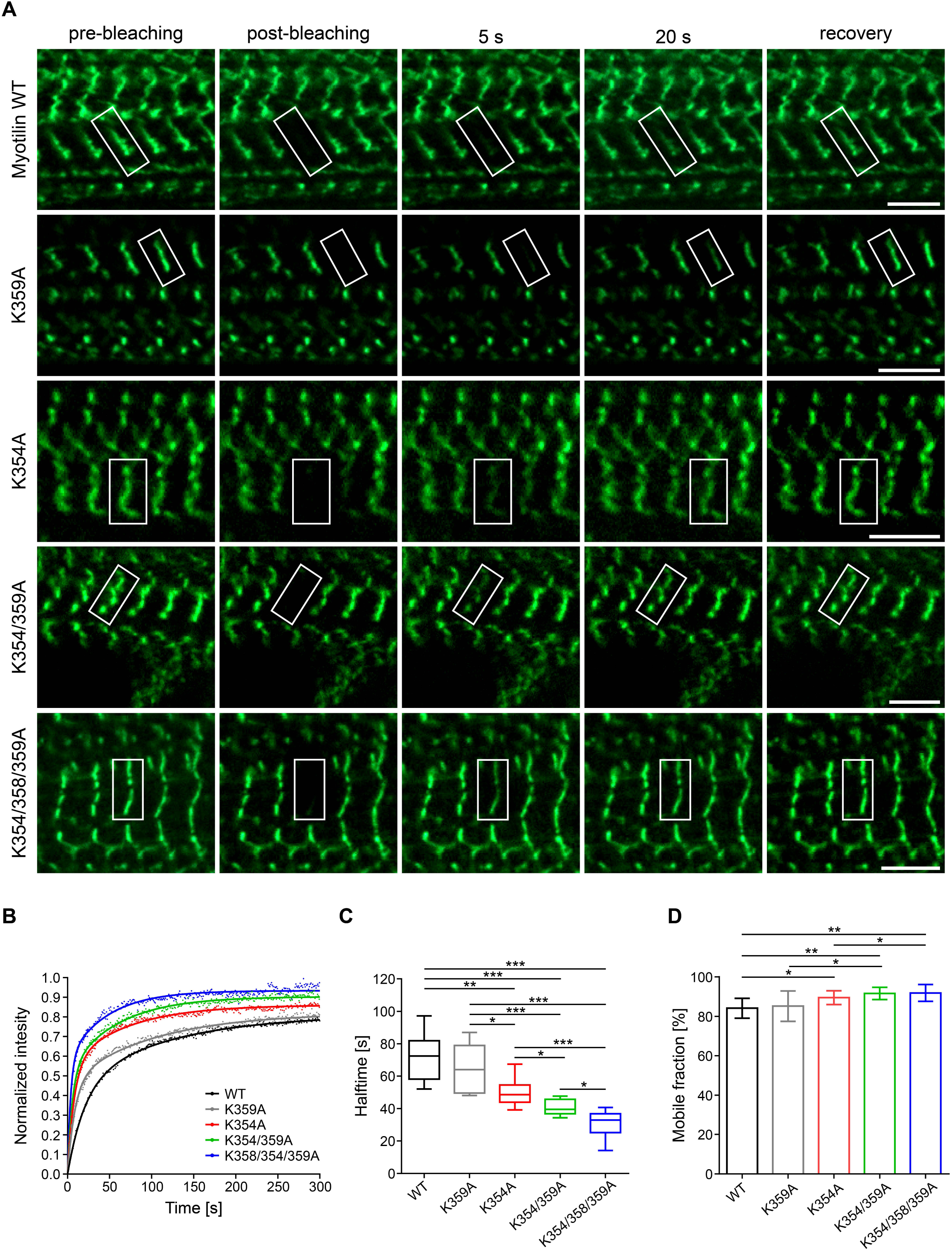
Binding of myotilin with F-actin affects its mobility and dynamics. (**A**) FRAP analysis of the dynamics of myotilin and its mutant variants in Z-discs. Localization of EGFP-fusion proteins of wild-type (WT) myotilin and its variants with the indicated mutations in Ig2 before bleaching (pre-bleaching), directly after bleaching (post-bleaching) and after the indicated time points after bleaching. White rectangles show regions of interePst (ROI), used for bleaching during FRAP experiments. Scale bars, 5 µm. (**B**) Representative FRAP recovery curves of the myotilin variants. (**C**) Halftimes of myotilin mutants compared to wild-type myotilin (WT). Quantification of data from FRAP studies shown in (**A** and **B**). Statistical data are depicted in box and whisker plots. Calculated median halftimes are shown as a line surrounded by a box, representing the interquartile range comprising the median ± 25% of the data. Whiskers extend at most two standard deviations from the median. (**D**) Mobile fractions of mutants compared to wild-type myotilin. Graph reports mean ± SEM, *n* = 8 myotubes for each variant. 1-3 Z-discs were bleached per myotube. Significance was assessed using 2-tailed Student’s *t*-test, *p < 0.1, **p < 0.05 and ***p < 0.01.

These results show that mutant myotilin variants with decreased binding to F-actin *in vitro* have significantly altered dynamics in the sarcomeric Z-disc, indicating that binding to F-actin stabilizes myotilin in the sarcomere.

## Discussion

Biochemical and structural analysis of myotilin and its interactions presented in this study provide a basis for the mechanistic understanding of its structural role in the sarcomeric Z-disc as a protein-protein interaction platform, and inferring its role in orchestration of Z-disc assembly and regulation of sarcomere biogenesis as an distinctive novel function of myotilin.

Our studies hint on Ig2 as the central F-actin interaction region, predominantly *via* electrostatic interactions. In addition they indicate that tandem Ig1Ig2 is the minimal functional F-actin-binding module, which the N- and C-terminal flanking regions further enhance binding affinity and thus contribute to complex stability. This is supported by a body of evidence: i) positively charged patches are on the same surface on both Ig domains, ii) XL-MS maps specific cross-links between F-actin and Ig1, Ig2 as well as the flanking regions, iii) the flanking regions enhance binding to F-actin, iv) NMR substantiated the interaction sites to map to the same face on both Ig domains, v) binding of tandem Ig1Ig2 to F-actin stabilizes F-actin and prevents its depolymerization. The latter may be explained by binding of each Ig domain to one actin subunit, allowing the tandem Ig1Ig2 to bridge two actin subunits and thus act as a vertical cross-linker. The importance of tandem Ig-domains for interaction with F-actin was also observed for palladin where Ig3Ig4 tandem displayed enhanced binding in comparison to Ig3 alone [18]. In addition, our FRAP experiments highlight the importance of the positively charged patch in Ig2 for binding. Proteins with charge-neutralizing mutations in Ig2 displayed increased mobile fractions and shorter fluorescence recovery times in differentiated C2C12 cells, indicating reduced binding affinity.

von Nandelstadh et al (2005) previously suggested Ig1Ig2 tandem to be important for optimal interaction with F-actin and showed that the size of myotilin constructs correlates with binding efficiency with longer fragments binding better to F-actin [17]. However, a detailed interaction map and molecular mechanism of the binding was not known. Based on our findings we generated the first molecular model of F-actin decorated by tandem Ig domains, which together with our quantitative binding assays allowed us to propose a mechanism for myotilin:F-actin assembly. In the unbound state, myotilin Ig domains are flexible, tending towards a semi-extended orientation which allows conformational plasticity as part of the binding-partner recognition mechanism. In the initial step of the interaction with F-actin, the positively charged region on myotilin Ig2 as the predominant binder, attaches to the negatively charged subdomain 1 of actin. The subsequent Ig1:F-actin interaction enables a transition to complete extension and consequently stabilization of both Ig domains and the flanking regions on F-actin.

Although palladin and myotilin are closely related, and the interaction with F-actin is in both cases mediated by a conserved mechanism involving basic charged clusters in the Ig-domains, they seem to differ fundamentally in the precise geometry and stoichiometry of the complexes formed. Palladin binds actin *via* two basic clusters situated on two sides of a single Ig-domain [28], allowing for cross-linking two actin filaments with a single palladin molecule. By contrast, a single myotilin molecule harbours only one high affinity binding site. Therefore, a dimerisation is required to enable F-actin cross-linking. Furthermore, the contribution of Ig-domain-flanking regions to F-actin binding may allow for regulation by post-translational mechanisms (PTMs) [4, 5].

Since the interaction of most actin-binding proteins with F-actin may either be regulated by certain compounds or modulated by other proteins, it is essential to contemplate the precise situation in the cell. For myotilin, it is therefore important to investigate potential roles of PIPs and known binding partners like α-actinin-2. α-Actinin-2 recognises the 1-4-5-8 motif of the cognate α-actinin-2 “neck” and/or titin Zr-7 by the pseudoligand mechanism regulated by PI(4,5)P_2_. Our studies suggest that myotilin binds to α-actinin-2 CAMD employing the same regulatory mechanism, setting the molecular basis of interaction for the entire palladin family. In addition, our binding data suggest that titin can displace myotilin from α-actinin-2, as titin Zr-7 interacts with α-actinin-2 with a stronger affinity (0.48 ± 0.13 µM) [7] than myotilin.

Furthermore, binding of PI(4,5)P_2_ to α-actinin-2 downregulates its F-actin binding activity, resulting in decreased F-actin cross-linking [50]. In palladin PI(4,5)P_2_ inhibits palladin-mediated F-actin polymerization and cross-linking [46]. By contrast, we showed that myotilin Ig1Ig2 does not bind PI(4,5)P_2_, however interactions with other parts cannot be completely excluded, suggesting that, as opposed to α-actinin-2 and palladin, the myotilin:F-actin interaction is not regulated by PI(4,5)P_2_. This implies a possible mechanism for stabilization of α-actinin-2 on F-actin upon binding to myotilin: PI(4,5)P_2_ binding induces α-actinin-2 open conformation enabling it to interact with myotilin, while its F-actin binding is simultaneously reduced due to PI(4,5)P_2_. Concurrent binding of myotilin to α-actinin-2 and F-actin can thus help to stabilize α-actinin-2 on F-actin and may compensate for the inhibitory effect of PI(4,5)P_2_.

Our SAXS data revealed full-length myotilin as a dynamic ensemble of multiple conformations with local structurally compact clusters, as previously predicted based on sequence analysis [22]. Further local induction of structure might be promoted upon binding, in particular to α-actinin-2, similar to titin and palladin where the interaction with CAMD of α-actinin-2 induces folding of the binding motif into an α-helical structure [7, 43, 45].

Our studies of myotilin dimerization in solution revealed that the regions flanking Ig1Ig2 domains enhance dimer stability. In comparison to the previous studies addressing myotilin dimerization [9, 10, 17], we show that myotilin dimerizes in a concentration-dependent manner and that only concentrations higher than 1.0 mM showed complete dimer population, implying a weak association. This suggests mechanisms driving or stabilizing dimer formation *in vivo*: high local concentrations and avidity effects mediated by concominat binding to interaction partners such as α-actinin, filamin C and of course F-actin that teether myotilin to the Z-disc, and/or PTMs.

All known pathogenic myotilin variants result in a single residue substitution in the protein, and all, except R405K, map to the N-terminus, within or close to the serine-rich region (SRR) and the hydrophobic residues-rich region (HRR) representing its “mutational hotspot” (**Figure 1B**). These mutations are predominantly substitutions from a polar/charged to a hydrophobic residue. Consequently, in concert with the slower degradation and changed turnover as already indicated previously [22, 24], a newly exposed hydrophobic cluster could promote aggregation and play an important role in pathological mechanisms. The R405K mutation in Ig2 was suggested to be responsible for defective homo-dimerization of myotilin and decreased interaction with α-actinin-2 [27]. However, our results revealed that R405K mutation does not impair myotilin homo-dimerization *in vitro* (**Figure S1G**), nor does it affect the binding of myotilin to F-actin (**Figure 2A**), which was expected since a positively charged residue is retained at this position. In addition, we predict this mutation will not have an impact on binding of myotilin to α-actinin-2, since it does not reside in the α-actinin-2 binding site or its proximity. Thus, further *in vitro* and *in vivo* experiments are needed to understand the molecular mechanism underlying the pathophysiology caused by the R405K mutation.

An intriguing yet unresolved question is: how are the multitude of F-actin-binding proteins sorted along thin myofilaments and Z-discs? Strikingly, tropomyosin is distributed all along thin filaments except the Z-disc region [6]. In non-muscle cells tropomyosin competes with α-actinin-1 and α-actinin-4, for F-actin binding, resulting in mutually exclusive localizations along stress fibers [34, 35]. We reveal that striated muscle-specific α-actinin-2 also competes with muscle type tropomyosin for binding to F-actin, and thus extend the competition mechanism to all isoforms of α-actinin. Most importantly, we found that myotilin also competes with tropomyosin for F-actin binding, and stabilizes F-actin against depolymerization, suggesting it may be involved in excluding tropomyosin from the Z-disc. Thus, binding of myotilin may create functionally distinct regions on actin filaments, thereby modulating the interactions of other proteins with F-actin. However, myotilin alone does not explain the absence of tropomyosin from the Z-disc. Instead, a concerted action of several Z-disc F-actin-binding proteins (e.g. filamin C, α-actinin-2, ZASP, myotilin etc.) is required, which is supported by the finding that the tropomyosin binding site on F-actin partly overlaps with the binding sites for filamin C, α-actinin-2 and myotilin (**Figure S4C**). Sorting will be facilitated by local protein complex formation, resulting in increased F-actin affinity, based on avidity effects. In this context it is noteworthy that even a dramatic reduction of the binding strength of myotilin to F-actin, as shown by our FRAP experiments, does not impair its Z-disc localization (**Figure 5**), supporting the view that the Z-disc is an intricate network of interacting multidomain proteins, which may tolerate reduction or even loss of individual interactions without attenuating the entire structure. This robustness rooted in redundancy may also explain that the loss of a single protein by gene knockout may be compensated by other proteins [51]. Accordingly, the lack of a phenotype in myotilin knockout mice may be compensated by myopalladin and/or palladin, which interact with EF34 hands of α-actinin-2 with similar affinity as myotilin [45].

This raises the question of the role of myotilin in Z-disc biogenesis. According to current models, in early myofibrillogenesis, premyofibrils are composed of minisarcomeres containing a subset of sarcomeric proteins in α-actinin-2 enriched Z-bodies, whereas the attached thin filaments are associated with tropomyosin and troponin [52]. A recent study showed that tropomyosins are present in sufficient levels in the cell to saturate all actin filaments [53]. Therefore tropomyosin needs to be actively inhibited from binding specific regions of the actin filaments to allow the formation of tropomyosin-free actin filaments [53] necessary for the development of Z-discs. Myotilin is expressed at relatively late stages of differentiation, when α-actinin-2, the Z-disc portion of titin, myopodin/SYNPO2 and filamin C already co-localize at Z-bodies [10, 54, 55]. Myotilin entry into this still relatively loose protein assembly occurs at the time when mechanical strain increases due to incipient contractile activity. Its simultaneous binding to F-actin and α-actinin, as well as the PIP_2_-dependent Zr7:α-actinin-2 interaction and the simultaneous displacement of tropomyosin make myotilin an ideal co-organizer of Z-disc proteins (**Figure 6**). This timing seems essential since the longer α-actinin molecules organize the more widely spaced F-actin networks required for myofibrils, whereas myotilin alone would form too tightly packed filament bundles. Further studies are required to determine the precise sequence of events regarding the assembly of components in maturing Z-discs.

**Figure 6.**
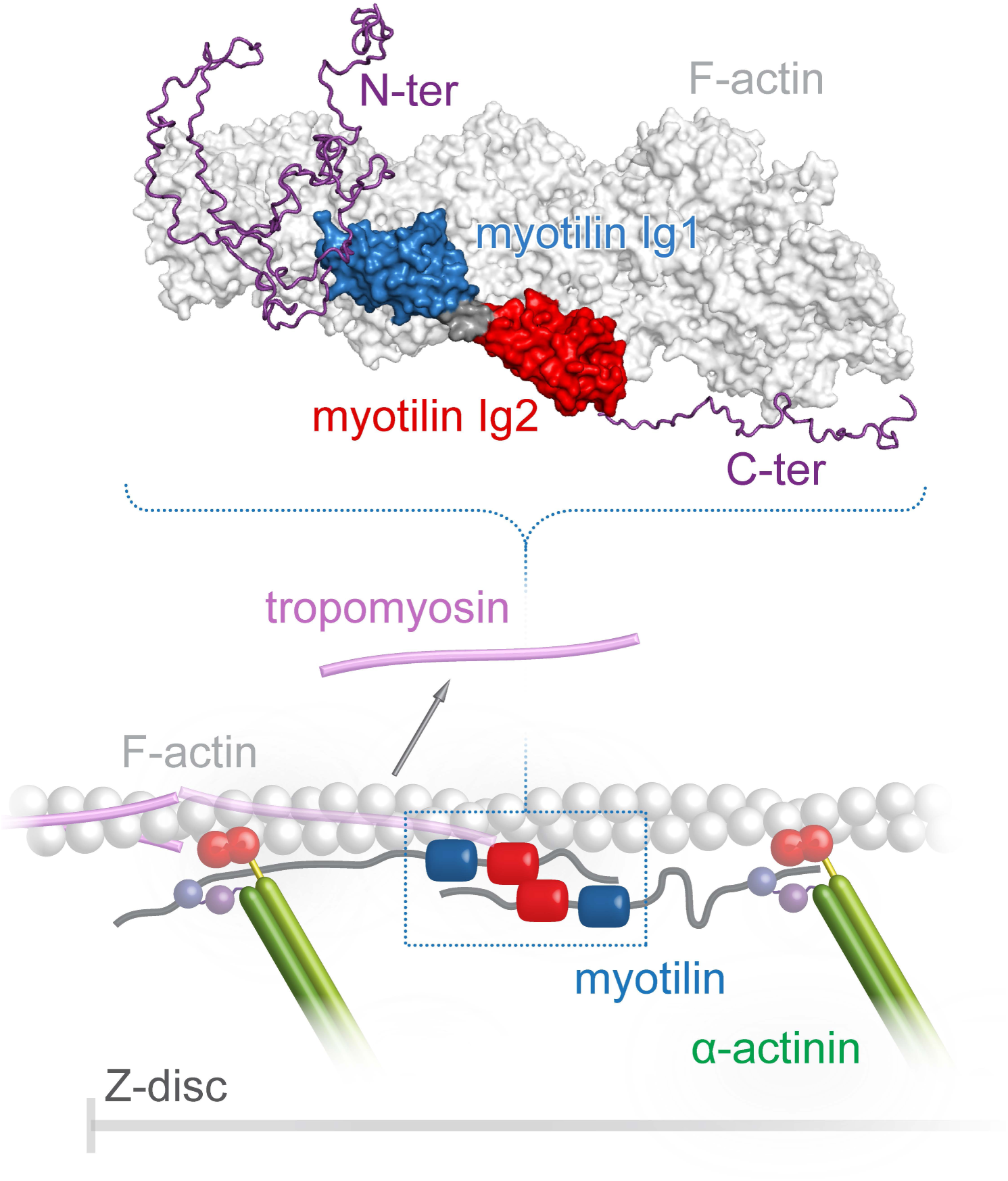
Model illustrating the role of myotilin in the organization of sarcomeric Z-disc assembly. Myotilin is recruited to the Z-disc by binding to F-actin, α-actinin-2 and/or other Z-disc components e.g. filamin C, ZASP. As a part of the recruitment process, myotilin can further stabilise (in its proposed antiparallel dimeric form [10] two α-actinin፧2 dimers on F-actin through interaction of its N-terminal region with CAMD of α-actinin፧2. Consequently, together with α-actinin-2 and/or other Z-disc components, myotilin can help to displace tropomyosin from the Z-disc. For simplicity, only some of the Z-disc components are shown.

## Materials and methods

### Plasmids and DNA constructs

Different DNA fragments, encoding human myotilin, or α-actinin-2 were sub-cloned into different vectors as specified in **Table S3**. For preparation of myotilin mutants (**Table S3**), mutagenesis was performed with the QuikChange II Site-Directed Mutagenesis Kit (Agilent). Trx-MYOT, MYOT^WT^ and Ig1Ig2^250-444^ constructs were used as templates in the PCR reactions. Correct assembly of the prepared constructs and mutagenesis efficiency was verified by DNA sequencing (GATC Biotech).

### Protein expression and purification

All recombinant myotilin and α-actinin-2 constructs were expressed in *Escherichia coli* (see **Table S3**). Full-length myotilin (Trx-MYOT) with N-terminal thioredoxin tag (Trx) and its mutant variant Trx-MYOT-NEECK were expressed in B834 (DE3) grown at 37°C in LB media to an OD_600_ of approximately 1.0. Protein expression was induced by addition of IPTG to a final concentration of 0.5 mM and cells were grown for additional 4-5 h at 37°C. Proteins were further purified by affinity chromatography using HisTrap FF columns (Cytiva) followed by size exclusion chromatography (SEC) on a HiLoad 26/600 Superdex 200 column (Cytiva) equilibrated with 20 mM Tris, 400 mM NaCl, 250 mM arginine, 2 mM β-mercaptoethanol, pH 7.5. Ig1Ig2^185-498^ was expressed in C41 (DE3) strain following the protocol described in [7] and purified by affinity chromatography using HisTrap FF columns. Subsequently, the Trx-tag was cleaved overnight with HRV-3C protease added at a mass ratio protease:protein of 1:1000, and the Ig1Ig2^185-498^ was further purified by StrepTrap HP columns (Cytiva), followed by SEC on a HiLoad 26/600 Superdex 75 column (Cytiva) equilibrated with 20 mM Tris, 150 mM NaCl, 5% glycerol, pH 7.5.

During purification of Ig1Ig2^185-498^*, the protein readily degrades from its C-terminus and forms Ig1Ig2^185-454^ as determined by intact protein mass spectrometry analysis of the final product. Thus to prepare Ig1Ig2^185፧454^, Ig1Ig2^185-498^* construct was expressed as described elsewhere [56]. After expression, cells were lysed by sonication and insoluble material removed by centrifugation. The lysate was loaded onto a GSTrap FF column (Cytiva) equilibrated with 10 mM sodium phosphate, 140 mM NaCl, 2.7 mM KCl, pH 7.4 (PBS). After washing with the same buffer the bound proteins were eluted by a linear gradient of 50 mM Tris, 10 mM reduced glutathione, pH 8.0. His_6_-GST-tag was removed by incubation with His_6_-tagged TEV protease (added at a mass ratio protease:protein of 1:100) during overnight dialysis against the dialysis buffer (50 mM Tris, 150 mM NaCl, 1 mM DTT, pH 7.4) at 4°C. The His_6_ -GST-tag-free Ig1Ig2^185-454^ was recovered as a flow-through after applying the cleavage mixture onto Ni^2+^-loaded HiTrap IMAC FF (Cytiva) and GSTrap FF columns to remove TEV protease, uncleaved proteins, and the cleaved-off His_6_-GST moiety. Final purification step was SEC on a Superdex 75 10/300 column (Cytiva) equilibrated with 20 mM HEPES, 150 mM NaCl, 5% glycerol, 1 mM DTT, pH 7.4.

During expression, Ig1Ig2^250-498^ degrades to Ig1Ig2^250-466^ as verified by intact protein mass spectrometry analysis. Thus, depending on the expression/purification strategy either Ig1Ig2^250-498^ or Ig1Ig2^250-466^ can be prepared from cells expressing Ig1Ig2^250-498^. To obtain Ig1Ig2^250-466^ the expression and purification procedure is basically the same as in [22] with the following differences: (1) for lysis buffer 20 mM HEPES, pH 7.4, 150 mM NaCl, 5% glycerol, 10 mM imidazole was used, the lysate was applied to HisTrap column and for elution the same buffer supplemented with 180 mM imidazole was used; (2) cleavage with HRV-3C protease was done at protease:protein mass ratio 1:50 using 20 mM HEPES, 150 mM NaCl, 5% glycerol, pH 7.4; (3) the final SEC step was done using a HiLoad 16/600 Superdex 75 column (Cytiva) equilibrated with 20 mM HEPES, 150 mM NaCl, 5% glycerol, pH 7.2. This same procedure is used to prepare Ig1Ig2^220፧452^, and Ig1Ig2^220፧452 R405K^. To obtain Ig1Ig2^250-498^, the procedure was the same as for Ig1Ig2^250-466^, however protein expression was done in B834 (DE3) strain at 37°C for 3 h after induction with 0.5mM IPTG. In addition, before final SEC step, Ig1Ig2^250-498^ was purified by cation exchange chromatography using Resource S column (Cytiva) in 10 mM HEPES, 2 M urea, 5% glycerol and gradient of 0-1 M NaCl, pH 7.0.

The Ig1Ig2^250-444^ and all its mutant variants (see **Table S3**) were purified as described elsewhere [22]. The Ig1^250-344^ and Ig2^349-459^ constructs were purified as described in [56]; the only modification was in the final SEC step where the buffer was replaced by 20 mM HEPES, 150 mM, NaCl, 5% glycerol, 1 mM DTT, pH 7.4.

Full-length α-actinin-2 constructs (ACTN2-WT and ACTN2-NEECK) were prepared as described previously [7]. ACTN2-EF14 was expressed as described in [7] and purified by affinity chromatography using HisTrap FF columns. Subsequently, the His_6_-tag was removed by incubation with HRV-3C protease added at a mass ratio protease:protein of 1:50 during overnight dialysis against PBS, 5% glycerol, 1 mM EDTA, 1 mM DTT at 4°C. The His_6_-tag-free ACTN2-EF14 was recovered as a flow-through after applying the cleavage mixture onto HisTrap FF column to remove HRV-3C protease, and uncleaved proteins. Final purification step was SEC on a HiLoad 26/600 Superdex 75 column equilibrated with PBS, 5% glycerol.

Palladin Ig3 was kindly provided by Moriah R. Beck (Wichita State University, Wichita, USA) and prepared/purified as described in [46]. Plasmid encoding constitutively monomeric mutant of actin (DVD-actin) was kindly provided by Michael K. Rosen (University of Texas Southwestern Medical Center, Dallas, USA) and protein was prepared as described in [30]. Doc2b [47] and tropomyosin [33] were kindly provided by Sascha Martens (Max Perutz Labs, University of Vienna, Vienna, Austria) and Stefan Raunser (Max Planck Institute of Molecular Physiology, Dortmund, Germany), respectively. Actin was prepared from rabbit skeletal muscle [57] and pyrene labeled following [58].

### SAXS data collection and analysis

SEC-SAXS data collection of Trx-MYOT in 20 mM Tris, 400 mM NaCl, 250 mM arginine, 5% glycerol, pH 7.5 was conducted at the EMBL P12 beamline of the storage ring PETRA III (DESY, Hamburg, Germany) [59] using an incident beam size of 200፧× ፧110፧μm^2^ (FWHM) in a 1.7፧mm quartz capillary held under vacuum. For this, the set-up as described in [60] was employed. Here, an integrated micro splitter (P-451, Upchurch Scientific) allows the eluent of a chromatography column (Superdex 200 increase 10/300 (Cytiva)) to flow equally through the SAXS capillary and a modular triple detector array (TDA, Viscotek model TDA 305, Malvern Panalytical) that extracts MW estimates of SEC-separated components by correlating refractive index (RI) and/or UV-vis concentrations with right angle laser light data (RALLS) using the integrated Omnisec software (Malvern Panalytical). In parallel, 3000 individual SAXS frames were collected with 1፧s exposure that were used for subsequent SAXS analysis. The chromatography was conducted at 0.5 ml/min. SAXS data reduction to produce the final scattering profile of monomeric Trx-MYOT was performed using standard methods. Briefly, 2D-to-1D radial averaging was performed using the SASFLOW pipeline [61]. Aided by the integrated prediction algorithms in CHROMIXS, the optimal frames within the elution peak and the buffer regions were selected [62]. Subsequently, single buffer frames were subtracted from sample frames one by one, scaled and averaged to produce the final subtracted curve which was further processed with various programs from the ATSAS software package [61]. Details on data collection are shown in **Table S1**.

SAXS data for purified myotilin constructs Ig1Ig2^250-444^ and Ig1Ig2^250-498^ was collected at ESRF beamline BM29 BioSAXS (Grenoble, France) equipped with the Pilatus 1M detector. Samples were measured at concentrations up to 14.12 (Ig1Ig2^250-444^) or 42.8 mg/ml (Ig1Ig2^250-498^) in 20 mM HEPES, 150 mM, NaCl, 5% glycerol, 1 mM DTT, pH 7.4, in two independent measurements. One SAXS dataset for the purified myotilin construct Ig1Ig2^220-452^ was collected at the EMBL X33 beamline of the storage ring Doris (DESY, Hamburg, Germany) equipped with the Pilatus 1M-W detector at concentrations in the range from 1 to 52 mg/ml in 20 mM MES, 200 mM NaCl, 3 % glycerol, pH 6.0. The other SAXS dataset for the myotilin construct Ig1Ig2^220-452^ and the Ig1Ig2^220-452 R405K^ mutant was collected at EMBL P12 beamline of the storage ring Petra III (DESY, Hamburg, Germany) equipped with the Pilatus 6M detector in the concentration range 2.1-36.6 mg/ml (Ig1Ig2^220-452^) and 2.3-45.5 mg/ml (Ig1Ig2^220-452 R405K^) in 20 mM HEPES, 150 mM NaCl, 5 % glycerol, 1 mM DTT, pH 7.4. The momentum of transfer *s* is defined as *s*=4πsinq/l. Details on data collection are shown in **Table S1**. For all constructs background scattering was subtracted, data reduced, normalised according to the measured concentration, and extrapolated to infinite dilution using the two lowest measured concentrations using PRIMUS [63] module of the ATSAS software package [61]. Forward scattering (I_0_) and radius of gyration (R_g_) were obtained by fitting the linear Guinier region of the data. Pair distribution function P(r) with the corresponding maximum particle size parameter (D_max_) was determined using GNOM program [64].

For reconstruction of a theoretical molecular envelope, *ab initio* modelling was performed 20 times using the program DAMMIF [65], where scattering from the calculated envelopes was fitted against the experimental scattering and evaluated by the chi values. The most typical envelope was selected by comparing the normalised spatial discrepancy (NSD) values between pairs of envelopes and later averaged by DAMAVER set of programs [66]. In rigid body modelling, high resolution structures of myotilin Ig1 (PDB: 2KDG) and Ig2 (PDB: 2KKQ) were used to fit SAXS scattering data using program CORAL [67]. Linker residues were designated as dummy atoms. *Ab initio* calculated envelope was superposed to the rigid body model using SUPCOMB program [68].

Flexibility analysis for Trx-MYOT was performed using EOM 2.1 program [26] with enforced P1 symmetry while the linker between Ig domains was again designated as dummy atoms. To account for the available information about the structure, the ensembles were generated as follows: during the generation, the potentially flexible termini/domain were allowed to have randomized conformations. Conformations consistent with scattering data were selected from the pool of 50 000 models using a genetic algorithm. The flexibility analysis was independently repeated three times; all runs gave comparable results. One should, however, note that, for flexible macromolecules exploring a range of conformations in solution, shape restoration returns average over the ensemble at a low resolution of the model in **Figure 1D** of 56+/-4 Å. All structure figures were prepared using PyMOL (The PyMOL Molecular Graphics System, version 2.3, Schrödinger, LLC). The SAXS data was deposited at SASBDB, accession: SASDFZ7, SASDF28, SASDF38, SASDF48.

### Cross-linking and mass spectrometry analyses

For cross-linking experiments with F-actin, actin was let to polymerize in PBS for 30 min at room temperature before adding 4-(4,6-dimethoxy-1,3,5-triazin-2-yl)-4-methylmorpholiniumchloride (DMTMM, MERCK) cross-linker. After incubation of F-actin with DMTMM for 5 min at room temperature Ig1Ig2^250፧444^, or Ig1Ig2^185፧454^, or Ig1Ig2^185፧498^ in PBS was added at concentrations indicated on the **Figure S3** and proteins were cross-linked for additional 40 min at room temperature. While for cross-linking of Ig1Ig2^250፧444^, increasing concentrations of DMTMM were used (**Figure S3A**), fixed concentration of DMTMM (2 mM) was used to cross-link Ig1Ig2^185፧454^ and Ig1Ig2^185፧498^ with F-actin (**Figures S3B and S3C**). Following separation of cross-linking samples by SDS-PAGE and staining of proteins with colloidal Coomassie Blue G-250, protein bands were excised, subjected to in-gel digestion using trypsin and analyzed by HPLC–ESI–MS/MS using an UltiMate 3000 RSLCnano system coupled to a Q Exactive Plus (both Thermo Fisher Scientific) mass spectrometer as described [69]. For identification of cross-linked peptides, the software pLink [70] was used, version 1.22 for Ig1Ig2^250-444^ and Ig1Ig2^185-454^, and version 2.0 for Ig1Ig2^185-498^. For the former, MS raw data files were first converted to the Mascot generic format (mgf) using msConvert from ProteoWizard, release 3.0.9740 [71]. Searches were performed against the forward and reversed amino acid sequences of the recombinant proteins with parameters and filtering criteria as described previously [72]. In brief, cross-linked peptide matches were filtered at a false discovery rate (FDR) of 5%, and a minimum E-value of 5 × 10^−2^ rejecting adjacent tryptic peptides of the same protein.

### Actin co-sedimentation assays

Actin co-sedimentation assays were performed as was published in [73], with some modifications. As various myotilin fragments displayed different solubility mostly depending on the pH of the buffer used within the assay, co-sedimentation assays were performed at two different conditions, B1 and B2. In the B1 conditions, rabbit skeletal muscle actin in G-buffer (2 mM HEPES, pH 8.0, 0.2 mM CaCl_2_, 0.2 mM ATP, 2 mM β-mercaptoethanol) was polymerized with the addition of 1/10 volume of 10x F-buffer (100 mM HEPES, pH 7.4, 500 mM KCl, 20 mM MgCl_2_) and incubated for 30 min at room temperature. Purified myotilin constructs Ig1^250-344^, Ig2^349-459^, Ig1Ig2^250-444^ and its mutant variants were dialyzed overnight against the dialysis buffer (20 mM HEPES, pH 7.4, 100 mM NaCl, 5% glycerol, 1 mM DTT). For apparent dissociation constant determination, fixed concentration (16 μM) of myotilin constructs in 20 μL of dialysis buffer was mixed with pre-polymerized F-actin (0-120 μM) in 20 μL of 1x F-buffer. For co-sedimentation with the increasing salt concentrations, pre-polymerized actin (6 μM) in 60 μL of 1x F-buffer was added to the myotilin construct in 60 μL of 1x F-buffer supplemented with or without KCl to achieve a final concentration of 50, 100, 150, 250 and 300 mM. For determination of binding of mutant myotilin variants to F-actin, pre-polymerized actin (8 μM) in 60 μL of 1x F-buffer were added to the mutant myotilin variants (8 μM) in 60 μL of dialysis buffer. In the B2 conditions, the G-buffer was 2 mM PIPES, pH 6.8, 0.2 mM CaCl_2_, 0.2 mM ATP, 5 mM β-mercaptoethanol, 10x F-buffer consisted of 100 mM PIPES, 1 M NaCl, 10 mM MgCl_2_, 10 mM EGTA, pH 6.8, and myotilin fragments Trx-MYOT, Ig1Ig2^250-444^, Ig1Ig2^250-498^, Ig1Ig2^185-454^, and Ig1Ig2^185-498^ were dialyzed overnight against the 10 mM PIPES, 100 mM NaCl, 1 mM MgCl_2_, 1 mM EGTA, 5% glycerol, pH 6.8. For determination of apparent dissociation constant(s) at B2 conditions, fixed concentration (6 μM) of myotilin fragment in 20 μL of dialysis buffer was mixed with pre-polymerized F-actin (0-60 μM) in 20 μL of 1x F-buffer. Co-sedimentation assays with tropomyosin and Trx-MYOT or ACTN2-NEECK mutant of α-actinin-2 were performed at B2 conditions as well. Here, Trx-MYOT (at final concentration 0-4.8 µM), or ACTN2-NEECK (0-6.0 µM) were mixed with pre-polymerized actin (6 µM) 30 min before the addition of tropomyosin (2 µM), or tropomyosin (2 µM) was mixed with pre-polymerized actin (6 µM) 30 min before addition of Trx-MYOT (at final concentration 0-4.8 µM). Reaction mixtures in all co-sedimentation assays were incubated at room temperature for 30 min prior to the ultracentrifugation (125.000 x g, 30 min, 22°C). Pellets and supernatants were separated and equal volumes analyzed by SDS-PAGE. Gels were scanned and quantified by densitometry with the QuantiScan 1.5 software (Biosoft). For determination of dissociation constants, the amount of bound myotilin constructs to F-actin in respect to free form was fitted with an equation for one site specific binding using Prism 6 (GraphPad Software).

### Preparation of liposomes and liposome co-sedimentation assay

Liposomes were prepared from chloroform stocks of 10 mg/ml of 1-palmitoyl-2-oleoyl-sn-glycero-3-phosphocholine, POPC, 16:0-18:1 (Avanti Polar Lipids) and 1 mg/ml of porcine brain L-α-phosphatidylinositol-4,5-bisphosphate, PI(4,5)P_2_ (Avanti Polar Lipids). Lipid mixtures of 0-20% of PI(4,5)P_2_ and 80-100% of POPC were first dried under a stream of nitrogen, and then vacuum dried for at least one hour. Dried lipid mixtures were hydrated in 20 mM Tris, 200 mM NaCl, 2% glycerol, 0.5 mM DTT, pH 8.5, incubated for 10 min at room temperature, gently mixed, further incubated for 20 min, and sonicated for 2 min at room temperature in a water bath, followed by 5 freezing-thawing cycles. Using a mini-extruder (Avanti Polar Lipids), vesicles were then extruded through polycarbonate filters with 0.4 µm and 0.1 µm pore sizes, respectively. In order to avoid the presence of potential aggregates, the Ig3 domain of palladin, Ig1Ig2^250-444^, and Doc2b, a positive control for PI(4,5)P_2_ binding, were first centrifuged in a TLA-55 rotor for 30 min at 100.000 x g, 4°C, using an optima MAX-XP benchtop ultracentrifuge (Beckman Coulter Life Sciences). For the assay, proteins and liposomes were mixed in polycarbonate tubes and incubated for 30 min at room temperature. Final concentrations of liposomes and proteins were 0.5 mg/ml and 10 µM, respectively. Mixtures were ultracentrifuged in TLA-100 rotor for 40 min at 100.000 x g, 20°C. Pellets and supernatants were separated and equal volumes analyzed by SDS-PAGE. Gels were scanned and quantified by densitometry with the QuantiScan 1.5 software (Biosoft).

### NMR spectroscopy

For all NMR-based measurements uniformly ^15^N-labelled constructs Ig1^250-344^, Ig2^349-459^ and Ig1Ig2^250-444^ were expressed in *E. coli* BL21(DE3) using standard M9 minimal media. To generate protein quantities amenable to ^1^H-^15^N HSQC based NMR studies, cell mass was generated in full media, before switching to minimal media for induction [74]. Protein purification was carried out as described above (see Protein expression and purification). Assignments for the Ig1 and Ig2 domains of myotilin were carried out previously and could be obtained from the Biological Magnetic Resonance Bank (Ig1, Entry Number: 7113; Ig2, Entry Number: 16370).

Assignments were reviewed in the original measurement conditions and were then transferred to the conditions used in this study by step gradients. For the construct Ig1Ig2^250-444^ no assignments were available and low expression yield was prohibitive to carrying out an assignment *de-novo*. Spectra of all three forms were therefore simply overlaid, which enabled us to generate a partial assignment of Ig1Ig2^250-444^. This was possible as the two domains seem to fold independently of each other and also do not show a dramatic change in shifts in the context of the full-length protein. A high affinity slowly exchanging interaction of myotilin and F-actin would lead to a combined molecular mass well above the NMR size limit. Weakly interacting species in fast exchange can, however, be measured on the free form, which retains favourable tumbling times and still yields information on the bound form, as the weighted average of the two species is detected. We thus used the shorter myotilin constructs displaying the weakest affinity to F-actin, as well as single domains.

The interaction of Ig1^250-344^, Ig2^349-459^, Ig1Ig2^250-444^ with F-actin was monitored using ^1^H-^15^N HSQC measurements [75, 76]. Various concentrations were used in these experiments and are indicated in the corresponding figure legends (**Figure S3D and S3E**). Both shift perturbations and changes in peak intensity due to the addition of F-actin were followed. While intensity changes were monitored as a simple ratio between the signal intensities (I_bound_/I_free_), shifts were tracked as a weighted average between proton and nitrogen shift changes in peak centres. All figures show the root of the sum of squared proton and nitrogen shifts in ppm values. While the proton shift is taken “as is”, the nitrogen shift is divided by a factor of five in order to reflect the fact that dispersion in the nitrogen dimension is higher, while the achieved resolution is lower. Due to these circumstances, a diagonal shift has about 5 times higher changes in the nitrogen versus the proton dimension 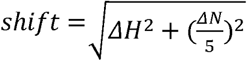. In globular domains composed of β-sheets like Ig-domains of myotilin, chemical shift changes are typically less localised and more dispersed on residues through the hydrogen-bonding network stabilising adjacent β-strands in the β-sheet. We thus did not focus on individual residues, but rather on segments of residues, which are part of the interaction interface. Shift averages showing shift-patterns were therefore calculated by application of a rolling window average over 11 residues where a minimum of 2 values were required to obtain a value for the average. The z-scores given express the difference of the observed value (x) from the mean (µ) in terms of multiples of the standard deviation (σ), i.e. 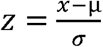. The average shift µ was calculated as a simple average 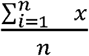.The standard deviation σ was calculated as 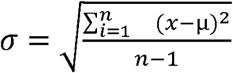, n being the number of all observed values.

NMR measurements were carried out on Bruker Avance spectrometer at 800 MHz and 298 K. Samples were prepared in 10 mM PIPES, 100 mM NaCl, 1 mM MgCl_2_, 5% glycerol, pH 6.8 and D_2_O was used at a final concentration of 10% as the lock solvent. The only exception is the F-actin gradient with Ig1, which was carried out in 10 mM imidazole, 50 mM KCl, 1 mM MgCl_2_, 1 mM EGTA, pH 7.1. Measurement at a single F-actin concentration at pH 6.8 confirms that the intensity changes and shifts induced by F-actin are very similar in both conditions. Spectra were processed using NMRPipe [77]. All spectra recorded used Rance-Kay detected sensitivity enhanced Heteronuclear Single Quantum Coherence sequences (HSQC) [75, 76].

### Macromolecular modeling

The Ig1Ig2^220-452^ models displaying hypothetical dimer architectures were prepared by manually moving two identical subunit copies where the Ig1 and Ig2 domains were in an extended relative orientation. The two subunits were oriented in a way to mimic Ig-Ig interactions, which would support dimer formation and to minimize steric clashes. These models were then used to calculate scattering profile using Crysol [78], and *P(r)* using GNOM [64], both part the ATSAS software package [61].

For the model of myotilin:F-actin complex the model of tandem Ig1Ig2 domains in semi-extended conformation based on SAXS data was used as a starting unbound structure [22]. Linker residues connecting myotilin Ig1 and Ig2 domains (residues 341-347), and N-terminal tail of actin (residues 3-6) were designated as flexible. As a starting F-actin structure an actin dimer was used (PDB 3J8I). The initial model was prepared by solvated docking implemented in Haddock 2.2 [31] using residues identified by chemical cross-linking and co-sedimentation assays as restraints. The best model with regard to Z-score was further refined using RosettaDock 2.3 [79] to yield the final myotilin:F-actin model.

### Microscale thermophoresis

Microscale thermophoresis measurements were performed on the Monolith NT.115 (NanoTemper Technologies) using fluorescently labelled DVD-actin. Purified DVD-actin was labelled using Monolith NT protein labelling kit RED-NHS (Amine Reactive) dye (NanoTemper Technologies) according to the manufacturer’s instructions. The labelling reaction was performed in the supplied labelling buffer with a concentration of 20 µM DVD-actin. The labelled DVD-actin was diluted to 80 nM with the reaction buffer: 20 mM HEPES, 100 mM KCl, 0.1 mM CaCl_2_, 0.2 mM ATP, 1 mM DTT, 5% glycerol, 0.2% Tween-20, 0.5 mg/mL BSA, pH 8.0. Solutions of concentrated myotilin constructs were serially diluted in 2:1 ratio, with the same reaction buffer. 5 µL of labelled DVD-actin was added to the 15 µL of unlabelled myotilin constructs. Reaction mixtures were incubated for 15 min at 25°C and approximately 5 µL of solution was loaded into Monolith NT Standard Capillaries (NanoTemper Technologies). Measurements were performed at 25°C using 20% LED and 20% IR-laser power with 5 s/30 s/5 s laser off/on/off times. The myotilin dependent change in thermophoresis was described with a Hill function to determine the apparent dissociation constant *K*_*d*_ of interaction from three independent measurements.

### F-actin depolymerization assays

Pyrene-labelled-F-actin filaments (30 μM, 25% pyrene labelled) were prepared by a 25 min incubation at room temperature in buffer A (10 mM HEPES, 50 mM KCl, 2 mM MgCl_2_, pH 8.0) in the presence or absence of different concentrations of Ig1Ig2^250-444^ (at molar ratios myotilin to actin as indicated on **Figure 2E**). Depolymerization of actin filaments was induced by 100x dilution with buffer A to a final concentration of 0.3 μM. Pre-cut pipet tips were used for all manipulations of actin filaments, and care was taken to avoid filament shearing. Pyrene fluorescence was monitored at room temperature over time at an excitation of 365 nm and emission of 388 nm in a Jasco FP-6300 fluorescence spectrophotometer.

### Transfection of myoblasts and fluorescence recovery after photobleaching

C2C12 cells were grown in proliferation medium [15% FCS, 100 U/ml penicillin, 100 µg/ml streptomycin, 2 mM non-essential amino acids and 1 mM sodium pyruvate, in Dulbecco’s modified Eagle’s medium (DMEM) with GlutaMAX]. Cells were trypsinized and transfected by nucleofection according to the recommendations of the manufacturer (Lonza, Cologne, Germany) with N-terminal EGFP-myotilin wild-type and mutants (K354A, K359A, K354/359A, K354/358/359A) (for details see **Table S3**). After transfection, cells were seeded on glass coverslips (WPI, Berlin, Germany) in proliferation medium. The medium was changed 24 h after transfection and cells were allowed to differentiate at 90% confluence by changing the medium to differentiation medium (2% horse serum, 100 U/ml penicillin, 100 µg/ml streptomycin, 2 mM non-essential amino acids and 1 mM sodium pyruvate, in DMEM with GlutaMAX). All media and supplements were from LifeTechnologies. Seven days later FRAP experiments were performed using a Cell Observer Spinning Disk Confocal Microscope (Carl Zeiss, Jena, Germany) equipped with an external 473 nm laser coupled via a scanner (UGA-40, Rapp OptoElectronic, Hamburg, Germany). Cells were continuously kept at 37°C and 5% CO_2_. Zen 2012 software was used for image processing. For FRAP analysis eight independent experiments were performed. Regions of interest (ROI) were limited to a single Z-disc and for each cell 1-3 ROI were chosen and bleached, for a pulse time of 1 ms with eight iterations, using the 473 nm laser (100 mW) with 100% intensity. A series of three images was taken before bleaching. Immediately after photobleaching, fluorescence recovery was monitored with an interval of 0.1 – 1 s until the signal was fully recovered (300 s). The ImageJ package Fiji was used to determine the fluorescence intensity of bleached and unbleached areas at each time point. Raw data were transformed into normalized FRAP curves as previously described [80].

### Isothermal titration calorimetry (ITC)

ITC experiments were performed at 25°C on a MicroCal PEAQ-ITC calorimeter (Malvern Panalytical). Protein samples were dialyzed against 20 mM Hepes, 400 mM NaCl, 250 mM arginine, 5% glycerol, pH 7.5. All solutions were degassed before the ITC experiments. Titrations consisted of sequential injections, in which 400 μM of ACTN2-EF14 was added to a sample cell containing 20 μM of either Trx-MYOT or Trx-MYOT-NEECK. The heats of dilution of ACTN2-EF14 into the buffer were determined separately and subtracted from the titration prior to analysis. All data were analyzed using MicroCal PEAQ-ITC analysis software (Malvern Panalytical).

### Pull-down assay

Before each experiment, proteins were centrifuged at 100,000 × g for 30 min, 4°C. Trx-MYOT (5 µM) was mixed with various α-actinin-2 constructs (2.5-7.5 µM, for details see **Figure 4F**) in 0.1 ml of buffer P (20 mM Tris-HCl, 100 mM NaCl, 10 mM imidazole, 5% glycerol, pH 7.5) and incubated for 30 min at room temperature. 25 µl of Co^2+^-loaded Chelating Sepharose beads (Cytiva) were added to each protein mixture, followed by incubation for 10 min at room temperature. After extensive washing with buffer P, proteins bound to the beads were eluted with 50 mM Tris, 500 mM NaCl, 500 mM imidazole, 7 M urea, pH 8.0 and analysed by SDS-PAGE.

### Differential scanning fluorimetry (DSF) assay

DSF was done following the protocol of [81] using a pH-screen described in [82]. Briefly, a master mix of protein/dye was prepared by mixing 420 µL of 0.5 mg/mL of Trx-MYOT with 2.1 µL 5000x SYPRO Orange (Thermo Fisher Scientific) and filled up with 20 mM Tris, 400 mM NaCl, 250 mM arginine, 2 mM β-mercaptoethanol, pH 7.5 to 525 µL. 5 µL of the protein/dye master mix was added to each well of 96-well PCR-plate containing 20 µL of the individual pH-screen solution (**Figure S2C**). The plate was sealed, tapped to mix and centrifuged for 30 s at 1500 x g. For the stability screen, the temperature was increased from 15°C to 95°C in 0.5°C (10 seconds hold time) increments. A fluorescence reading was done every 0.5°C. Data analysis was performed using the CFX Manager software (Bio-Rad) included with the RT-PCR machine.

### Data availability

The atomic coordinates have been deposited in the SASDB and are available with these links:

SASDFZ7

Full-length myotilin with N-terminally fused thioredoxin tag

https://www.sasbdb.org/data/SASDFZ7/4tm5723u5y/

SASDF28

Myotilin immunoglobulin domains Ig1Ig2 (220-452)

https://www.sasbdb.org/data/SASDF28/b0sfyy2m3k/

SASDF38

Myotilin immunoglobulin domains Ig1Ig2 (250-444)

https://www.sasbdb.org/data/SASDF38/knazhmxat5/

SASDF48

Myotilin immunoglobulin domains Ig1Ig2 (250-498)

https://www.sasbdb.org/data/SASDF48/w8mwgpml0d/

SASDJH8

Myotilin immunoglobulin domains Ig1Ig2 (220-452, wild-type) concentration series data

https://www.sasbdb.org/data/SASDJH8/6fkwyvfgcl/

SASDJJ8

Myotilin immunoglobulin domains Ig1Ig2 (220-452, R405K mutant) concentration series data

https://www.sasbdb.org/data/SASDJJ8/af8sxzaqhi/

All other relevant data are described in SI Appendix or are available upon request. Full methods can be found in SI Appendix, Materials and Methods.

## Author contributions

Conceptualization: KD-C, JK, BL and DOF; investigation: JK, MP, VP, TCS, FD, SM, MAG, CS, SS; methodology: JK, MP, VP, TCS, FD, SM, MAG, CS, SS, PFMV; visualization: JK, MP, TCS, MAG, PFMV; data curation: KD-C; supervision: KD-C, DOF, BW, RK, DIS, BL; validation: KD-C, DOF, BW, RK, PFMV, JK, DIS; resources: KD-C, DOF, BW, RK, DIS; funding acquisition: KD-C, DOF, BW, RK, BL; writing (original draft): JK, MP, VP, TCS, FD, SM, MAG, SS, PFMV, DOF, KD-C; writing (review and editing): all authors.

## Acknowledgements

We thank the staff of the SAXS beamline at ESRF in Grenoble and EMBL-Hamburg for their excellent support, especially Martin Schroer for his assistance with the SEC-SAXS/RALS set-up. This work was supported by the Slovenian Research Agency young researcher grant (No. 35337) and research program P1-0140. KDC research was supported by a Marie Curie Initial Training Network: MUZIC (N°238423), Austrian Science Fund (FWF) Projects I525, I1593, P22276, P19060 and W1221, Federal Ministry of Economy, Family and Youth through the initiative “Laura Bassi Centres of Expertise” funding the Centre of Optimized Structural Studies, N°253275, by the Wellcome Trust Collaborative Award (201543/Z/16), Austrian-Slovak Interreg Project B301 StruBioMol, COST action BM1405 - Non-globular proteins - from sequence to structure, function and application in molecular physiopathology (NGP-NET), WWTF (Vienna Science and Technology Fund) Chemical Biology project LS17-008, and by the University of Vienna.

This study was further supported by the Deutsche Forschungsgemeinschaft (DFG, German Research Foundation) FOR 1352 (projects P1, D.F.; and P4, B.W.) FOR 2743 (projects P6, D.F.; and P9, B.W.) Project ID 403222702/SFB 1381 (project Z1, B.W.) and Germany’s Excellence Strategy (CIBSS – EXC-2189 – Project ID 390939984, BW).

The FP7 WeNMR (project# 261572), H2020 West-Life (project# 675858) and the EOSC-hub (project #777536) European e-Infrastructure projects are acknowledged for the use of their web portals, which make use of the EGI infrastructure with the dedicated support of CESNET-MetaCloud, INFN-PADOVA, NCG-INGRID-PT, TW-NCHC, SURFsara and NIKHEF, and the additional support of the national GRID Initiatives of Belgium, France, Italy, Germany, the Netherlands, Poland, Portugal, Spain, UK, Taiwan and the US Open Science Grid.

S. Raunser (Max Planck Institute of Molecular Physiology, Dortmund, Germany), M. Beck (Wichita State University, USA), S. Martins (University of Vienna, AT) and M.K. Rosen (University of Texas Southwestern Medical Center, USA) are acknowledged for kindly providing clones and/or protein samples for human tropomyosin, Ig3 domain of human palladin, Doc2b and non-polymerizable *Drosophila melanogaster* DVD actin mutant, respectively. We thank Life Science Editors for editing assistance.

## References

1. Janssen I, Heymsfield SB, Wang ZM, Ross R. Skeletal muscle mass and distribution in 468 men and women aged 18-88 yr. J Appl Physiol (1985). 2000;89(1):81–8. Epub 2000/07/25. doi:10.1152/jappl.2000.89.1.81. PubMed PMID:10904038.

2. Ulbricht A, Eppler FJ, Tapia VE, van der Ven PF, Hampe N, Hersch N, et al. Cellular mechanotransduction relies on tension-induced and chaperone-assisted autophagy. Curr Biol. 2013;23(5):430–5. Epub 2013/02/26. doi:10.1016/j.cub.2013.01.064. PubMed PMID:23434281.

3. Frank D, Frey N. Cardiac Z-disc signaling network. J Biol Chem. 2011;286(12):9897–904. Epub 2011/01/25. doi:10.1074/jbc.R110.174268. PubMed PMID:21257757; PubMed Central PMCID:PMCPMC3060542.

4. Reimann L, Schwable AN, Fricke AL, Muhlhauser WWD, Leber Y, Lohanadan K, et al. Phosphoproteomics identifies dual-site phosphorylation in an extended basophilic motif regulating FILIP1-mediated degradation of filamin-C. Commun Biol. 2020;3(1):253. Epub 2020/05/24. doi:10.1038/s42003-020-0982-5. PubMed PMID:32444788.

5. Reimann L, Wiese H, Leber Y, Schwable AN, Fricke AL, Rohland A, et al. Myofibrillar Z-discs Are a Protein Phosphorylation Hot Spot with Protein Kinase C (PKCalpha) Modulating Protein Dynamics. Mol Cell Proteomics. 2017;16(3):346–67. doi:10.1074/mcp.M116.065425. PubMed PMID:28028127; PubMed Central PMCID:PMCPMC5340999.

6. Luther PK. The vertebrate muscle Z-disc: sarcomere anchor for structure and signalling. J Muscle Res Cell Motil. 2009;30(5-6):171–85. doi:10.1007/s10974-009-9189-6. PubMed PMID:19830582; PubMed Central PMCID:PMCPMC2799012.

7. Ribeiro Ede A, Jr., Pinotsis N, Ghisleni A, Salmazo A, Konarev PV, Kostan J, et al. The structure and regulation of human muscle alpha-actinin. Cell. 2014;159(6):1447–60. doi:10.1016/j.cell.2014.10.056. PubMed PMID:25433700; PubMed Central PMCID:PMCPMC4259493.

8. Young P, Gautel M. The interaction of titin and alpha-actinin is controlled by a phospholipid-regulated intramolecular pseudoligand mechanism. EMBO J. 2000;19(23):6331–40. doi:10.1093/emboj/19.23.6331. PubMed PMID:11101506; PubMed Central PMCID:PMCPMC305858.

9. Salmikangas P, Mykkanen OM, Gronholm M, Heiska L, Kere J, Carpen O. Myotilin, a novel sarcomeric protein with two Ig-like domains, is encoded by a candidate gene for limb-girdle muscular dystrophy. Hum Mol Genet. 1999;8(7):1329–36. doi:10.1093/hmg/8.7.1329. PubMed PMID:10369880.

10. Salmikangas P, van der Ven PF, Lalowski M, Taivainen A, Zhao F, Suila H, et al. Myotilin, the limb-girdle muscular dystrophy 1A (LGMD1A) protein, cross-links actin filaments and controls sarcomere assembly. Hum Mol Genet. 2003;12(2):189–203. Epub 2002/12/25. doi:10.1093/hmg/ddg020. PubMed PMID:12499399.

11. Hauser MA, Horrigan SK, Salmikangas P, Torian UM, Viles KD, Dancel R, et al. Myotilin is mutated in limb girdle muscular dystrophy 1A. Hum Mol Genet. 2000;9(14):2141–7. doi:10.1093/hmg/9.14.2141. PubMed PMID:10958653.

12. Gontier Y, Taivainen A, Fontao L, Sonnenberg A, van der Flier A, Carpen O, et al. The Z-disc proteins myotilin and FATZ-1 interact with each other and are connected to the sarcolemma via muscle-specific filamins. J Cell Sci. 2005;118(Pt 16):3739–49. doi:10.1242/jcs.02484. PubMed PMID:16076904.

13. von Nandelstadh P, Ismail M, Gardin C, Suila H, Zara I, Belgrano A, et al. A class III PDZ binding motif in the myotilin and FATZ families binds enigma family proteins: a common link for Z-disc myopathies. Mol Cell Biol. 2009;29(3):822–34. Epub 2008/12/03. doi:10.1128/MCB.01454-08. PubMed PMID:19047374; PubMed Central PMCID:PMCPMC2630697.

14. Ruparelia A, Vaz R, Bryson-Richardso R. Myofibrillar Myopathies and the Z-Disk Associated Proteins. In: Cseri J, editor. Skeletal Muscle -From Myogenesis to Clinical Relations Croatia: InTech; 2012. p. 317–58.

15. Selcen D, Engel AG. Mutations in myotilin cause myofibrillar myopathy. Neurology. 2004;62(8):1363–71. doi:10.1212/01.wnl.0000123576.74801.75.

16. van der Ven PF, Wiesner S, Salmikangas P, Auerbach D, Himmel M, Kempa S, et al. Indications for a novel muscular dystrophy pathway. gamma-filamin, the muscle-specific filamin isoform, interacts with myotilin. J Cell Biol. 2000;151(2):235–48. doi:10.1083/jcb.151.2.235. PubMed PMID:11038172; PubMed Central PMCID:PMCPMC2192634.

17. von Nandelstadh P, Gronholm M, Moza M, Lamberg A, Savilahti H, Carpen O. Actin-organising properties of the muscular dystrophy protein myotilin. Exp Cell Res. 2005;310(1):131–9. doi:10.1016/j.yexcr.2005.06.027. PubMed PMID:16122733.

18. Dixon RD, Arneman DK, Rachlin AS, Sundaresan NR, Costello MJ, Campbell SL, et al. Palladin is an actin cross-linking protein that uses immunoglobulin-like domains to bind filamentous actin. J Biol Chem. 2008;283(10):6222–31. doi:10.1074/jbc.M707694200. PubMed PMID:18180288.

19. Nakamura F, Osborn TM, Hartemink CA, Hartwig JH, Stossel TP. Structural basis of filamin A functions. J Cell Biol. 2007;179(5):1011–25. PubMed PMID: Medline:18056414.

20. Otey CA, Dixon R, Stack C, Goicoechea SM. Cytoplasmic Ig-domain proteins: cytoskeletal regulators with a role in human disease. Cell Motil Cytoskeleton. 2009;66(8):618–34. doi:10.1002/cm.20385. PubMed PMID:19466753; PubMed Central PMCID:PMCPMC2735333.

21. Otey CA, Rachlin A, Moza M, Arneman D, Carpen O. The palladin/myotilin/myopalladin family of actin-associated scaffolds. Int Rev Cytol. 2005;246:31–58. doi:10.1016/S0074-7696(05)46002-7. PubMed PMID:16164966.

22. Puz V, Pavsic M, Lenarcic B, Djinovic-Carugo K. Conformational plasticity and evolutionary analysis of the myotilin tandem Ig domains. ci Rep. 2017;7(1):3993. doi:10.1038/s41598-017-03323-6. PubMed PMID:28638118; PubMed Central PMCID:PMCPMC5479843.

23. Konrat R. The protein meta-structure: a novel concept for chemical and molecular biology. Cell Mol Life Sci. 2009;66(22):3625–39. doi:10.1007/s00018-009-0117-0. PubMed PMID:19690801.

24. von Nandelstadh P, Soliymani R, Baumann M, Carpen O. Analysis of myotilin turnover provides mechanistic insight into the role of myotilinopathy-causing mutations. Biochem J. 2011;436(1):113–21. doi:10.1042/BJ20101672. PubMed PMID:21361873.

25. Keduka E, Hayashi YK, Shalaby S, Mitsuhashi H, Noguchi S, Nonaka I, et al. In Vivo Characterization of Mutant Myotilins. The American Journal of Pathology. 2012;180(4):1570–80. doi:10.1016/j.ajpath.2011.12.040.

26. Tria G, Mertens HD, Kachala M, Svergun DI. Advanced ensemble modelling of flexible macromolecules using X-ray solution scattering. IUCrJ. 2015;2(Pt 2):207–17. doi:10.1107/S205225251500202X. PubMed PMID:25866658; PubMed Central PMCID:PMCPMC4392415.

27. Shalaby S, Mitsuhashi H, Matsuda C, Minami N, Noguchi S, Nonaka I, et al. Defective myotilin homodimerization caused by a novel mutation in MYOT exon 9 in the first Japanese limb girdle muscular dystrophy 1A patient. J Neuropathol Exp Neurol. 2009;68(6):701–7. doi:10.1097/NEN.0b013e3181a7f703. PubMed PMID:19458539.

28. Beck MR, Dixon RD, Goicoechea SM, Murphy GS, Brungardt JG, Beam MT, et al. Structure and function of palladin’s actin binding domain. J Mol Biol. 2013;425(18):3325–37. doi:10.1016/j.jmb.2013.06.016. PubMed PMID:23806659; PubMed Central PMCID:PMCPMC3759364.

29. Marks DS, Hopf TA, Sander C. Protein structure prediction from sequence variation. Nat Biotechnol. 2012;30(11):1072–80. doi:10.1038/nbt.2419. PubMed PMID:23138306; PubMed Central PMCID:PMCPMC4319528.

30. Zahm JA, Padrick SB, Chen Z, Pak CW, Yunus AA, Henry L, et al. The bacterial effector VopL organizes actin into filament-like structures. Cell. 2013;155(2):423–34. doi:10.1016/j.cell.2013.09.019. PubMed PMID:24120140; PubMed Central PMCID:PMCPMC4048032.

31. van Zundert GCP, Rodrigues J, Trellet M, Schmitz C, Kastritis PL, Karaca E, et al. The HADDOCK2.2 Web Server: User-Friendly Integrative Modeling of Biomolecular Complexes. J Mol Biol. 2016;428(4):720–5. Epub 2015/09/28. doi:10.1016/j.jmb.2015.09.014. PubMed PMID:26410586.

32. Heikkinen O, Permi P, Koskela H, Carpen O, Ylanne J, Kilpelainen I. Solution structure of the first immunoglobulin domain of human myotilin. J Biomol NMR. 2009;44(2):107–12. doi:10.1007/s10858-009-9320-4. PubMed PMID:19418025.

33. von der Ecken J, Muller M, Lehman W, Manstein DJ, Penczek PA, Raunser S. Structure of the F-actin-tropomyosin complex. Nature. 2015;519(7541):114–7. doi:10.1038/nature14033. PubMed PMID:25470062; PubMed Central PMCID:PMCPMC4477711.

34. Gateva G, Kremneva E, Reindl T, Kotila T, Kogan K, Gressin L, et al. Tropomyosin Isoforms Specify Functionally Distinct Actin Filament Populations In Vitro. Curr Biol. 2017;27(5):705–13. doi:10.1016/j.cub.2017.01.018. PubMed PMID:28216317; PubMed Central PMCID:PMCPMC5344678.

35. Kemp JP, Jr., Brieher WM. The actin filament bundling protein alpha-actinin-4 actually suppresses actin stress fibers by permitting actin turnover. J Biol Chem. 2018;293(37):14520–33. doi:10.1074/jbc.RA118.004345. PubMed PMID:30049798; PubMed Central PMCID:PMCPMC6139541.

36. Galkin VE, Orlova A, Salmazo A, Djinovic-Carugo K, Egelman EH. Opening of tandem calponin homology domains regulates their affinity for F-actin. Nat Struct Mol Biol. 2010;17(5):614–6. doi:10.1038/nsmb.1789. PubMed PMID:20383143; PubMed Central PMCID:PMCPMC2921939.

37. Avery AW, Fealey ME, Wang F, Orlova A, Thompson AR, Thomas DD, et al. Structural basis for high-affinity actin binding revealed by a beta-III-spectrin SCA5 missense mutation. Nat Commun. 2017;8(1):1350. doi:10.1038/s41467-017-01367-w. PubMed PMID:29116080; PubMed Central PMCID:PMCPMC5676748.

38. Iwamoto DV, Huehn A, Simon B, Huet-Calderwood C, Baldassarre M, Sindelar CV, et al. Structural basis of the filamin A actin-binding domain interaction with F-actin. Nat Struct Mol Biol. 2018;25(10):918–27. doi:10.1038/s41594-018-0128-3. PubMed PMID:30224736; PubMed Central PMCID:PMCPMC6173970.

39. Janco M, Bonello TT, Byun A, Coster AC, Lebhar H, Dedova I, et al. The impact of tropomyosins on actin filament assembly is isoform specific. Bioarchitecture. 2016;6(4):61–75. doi:10.1080/19490992.2016.1201619. PubMed PMID:27420374; PubMed Central PMCID:PMCPMC6085118.

40. Pollard TD. Actin and Actin-Binding Proteins. Cold Spring Harb Perspect Biol. 2016;8(8). doi:10.1101/cshperspect.a018226. PubMed PMID:26988969; PubMed Central PMCID:PMCPMC4968159.

41. Way M, Pope B, Weeds AG. Evidence for functional homology in the F-actin binding domains of gelsolin and alpha-actinin: implications for the requirements of severing and capping. J Cell Biol. 1992;119(4):835–42. Epub 1992/11/01. doi:10.1083/jcb.119.4.835. PubMed PMID:1331120; PubMed Central PMCID:PMCPMC2289707.

42. Leinweber B, Tang JX, Stafford WF, Chalovich JM. Calponin interaction with alpha-actinin-actin: evidence for a structural role for calponin. Biophys J. 1999;77(6):3208–17. Epub 1999/12/10. doi:10.1016/S0006-3495(99)77151-1. PubMed PMID:10585942; PubMed Central PMCID:PMCPMC1289132.

43. Atkinson RA, Joseph C, Kelly G, Muskett FW, Frenkiel TA, Nietlispach D, et al. Ca2+-independent binding of an EF-hand domain to a novel motif in the alpha-actinin-titin complex. Nat Struct Biol. 2001;8(10):853–7. doi:10.1038/nsb1001-853. PubMed PMID:11573089.

44. Ronty M, Taivainen A, Moza M, Otey CA, Carpen O. Molecular analysis of the interaction between palladin and alpha-actinin. FEBS Lett. 2004;566(1-3):30–4. Epub 2004/05/19. doi:10.1016/j.febslet.2004.04.006. PubMed PMID:15147863.

45. Beck MR, Otey CA, Campbell SL. Structural characterization of the interactions between palladin and alpha-actinin. J Mol Biol. 2011;413(3):712–25. doi:10.1016/j.jmb.2011.08.059. PubMed PMID:21925511; PubMed Central PMCID:PMCPMC3226707.

46. Yadav R, Vattepu R, Beck MR. Phosphoinositide Binding Inhibits Actin Crosslinking and Polymerization by Palladin. J Mol Biol. 2016;428(20):4031–47. doi:10.1016/j.jmb.2016.07.018. PubMed PMID:27487483; PubMed Central PMCID:PMCPMC5525146.

47. Groffen AJ, Martens S, Diez Arazola R, Cornelisse LN, Lozovaya N, de Jong AP, et al. Doc2b is a high-affinity Ca2+ sensor for spontaneous neurotransmitter release. Science. 2010;327(5973):1614–8. doi:10.1126/science.1183765. PubMed PMID:20150444; PubMed Central PMCID:PMCPMC2846320.

48. Wang J, Dube DK, Mittal B, Sanger JM, Sanger JW. Myotilin dynamics in cardiac and skeletal muscle cells. Cytoskeleton (Hoboken). 2011;68(12):661–70. doi:10.1002/cm.20542. PubMed PMID:22021208; PubMed Central PMCID:PMCPMC3240742.

49. Wang J, Shaner N, Mittal B, Zhou Q, Chen J, Sanger JM, et al. Dynamics of Z-band based proteins in developing skeletal muscle cells. Cell Motil Cytoskeleton. 2005;61(1):34–48. doi:10.1002/cm.20063. PubMed PMID:15810059; PubMed Central PMCID:PMCPMC1993831.

50. Fraley TS, Tran TC, Corgan AM, Nash CA, Hao J, Critchley DR, et al. Phosphoinositide binding inhibits alpha-actinin bundling activity. J Biol Chem. 2003;278(26):24039–45. doi:10.1074/jbc.M213288200. PubMed PMID:12716899.

51. Moza M, Mologni L, Trokovic R, Faulkner G, Partanen J, Carpen O. Targeted deletion of the muscular dystrophy gene myotilin does not perturb muscle structure or function in mice. Mol Cell Biol. 2007;27(1):244–52. doi:10.1128/MCB.00561-06. PubMed PMID:17074808; PubMed Central PMCID:PMCPMC1800670.

52. Sanger JW, Kang S, Siebrands CC, Freeman N, Du A, Wang J, et al. How to build a myofibril. J Muscle Res Cell Motil. 2005;26(6-8):343–54. doi:10.1007/s10974-005-9016-7. PubMed PMID:16465476.

53. Meiring JCM, Bryce NS, Wang Y, Taft MH, Manstein DJ, Liu Lau S, et al. Co-polymers of Actin and Tropomyosin Account for a Major Fraction of the Human Actin Cytoskeleton. Curr Biol. 2018;28(14):2331-7.e5. PubMed PMID:Medline:29983319.

54. Linnemann A, van der Ven PF, Vakeel P, Albinus B, Simonis D, Bendas G, et al. The sarcomeric Z-disc component myopodin is a multiadapter protein that interacts with filamin and alpha-actinin. Eur J Cell Biol. 2010;89(9):681–92. doi:10.1016/j.ejcb.2010.04.004. PubMed PMID:20554076.

55. Mologni L, Moza M, Lalowski MM, Carpen O. Characterization of mouse myotilin and its promoter. Biochem Biophys Res Commun. 2005;329(3):1001–9. doi:10.1016/j.bbrc.2005.02.074. PubMed PMID:15752755.

56. Drmota Prebil S, Slapsak U, Pavsic M, Ilc G, Puz V, de Almeida Ribeiro E, et al. Structure and calcium-binding studies of calmodulin-like domain of human non-muscle alpha-actinin-1. Sci Rep. 2016;6:27383. doi:10.1038/srep27383. PubMed PMID:27272015; PubMed Central PMCID:PMCPMC4895382.

57. Spudich JA, Watt S. The regulation of rabbit skeletal muscle contraction. I. Biochemical studies of the interaction of the tropomyosin-troponin complex with actin and the proteolytic fragments of myosin. J Biol Chem. 1971;246(15):4866–71. PubMed PMID:4254541.

58. Kouyama T, Mihashi K. Fluorimetry study of N-(1-pyrenyl)iodoacetamide-labelled F-actin. Local structural change of actin protomer both on polymerization and on binding of heavy meromyosin. Eur J Biochem. 1981;114(1):33–8. PubMed PMID:7011802.

59. Blanchet CE, Spilotros A, Schwemmer F, Graewert MA, Kikhney A, Jeffries CM, et al. Versatile sample environments and automation for biological solution X-ray scattering experiments at the P12 beamline (PETRA III, DESY). J Appl Crystallogr. 2015;48(Pt 2):431–43. doi:10.1107/S160057671500254X. PubMed PMID:25844078; PubMed Central PMCID:PMCPMC4379436.

60. Graewert MA, Franke D, Jeffries CM, Blanchet CE, Ruskule D, Kuhle K, et al. Automated pipeline for purification, biophysical and x-ray analysis of biomacromolecular solutions. Sci Rep. 2015;5:10734. doi:10.1038/srep10734. PubMed PMID:26030009; PubMed Central PMCID:PMCPMC5377070.

61. Franke D, Petoukhov MV, Konarev PV, Panjkovich A, Tuukkanen A, Mertens HDT, et al. ATSAS 2.8: a comprehensive data analysis suite for small-angle scattering from macromolecular solutions. J Appl Crystallogr. 2017;50(Pt 4):1212–25. doi:10.1107/S1600576717007786. PubMed PMID:28808438; PubMed Central PMCID:PMCPMC5541357.

62. Panjkovich A, Svergun DI. CHROMIXS: automatic and interactive analysis of chromatography-coupled small-angle X-ray scattering data. Bioinformatics. 2018;34(11):1944–6. doi:10.1093/bioinformatics/btx846. PubMed PMID: 29300836; PubMed Central PMCID:PMCPMC5972624.

63. Konarev PV, Volkov VV, Sokolova AV, Koch MHJ, Svergun DI. PRIMUS: a Windows PC-based system for small-angle scattering data analysis. Journal of Applied Crystallography. 2003;36:1277–82. doi:10.1107/S0021889803012779. PubMed PMID:WOS:000185178600026.

64. Svergun DI. Determination of the Regularization Parameter in Indirect-Transform Methods Using Perceptual Criteria. Journal of Applied Crystallography. 1992;25:495–503. doi:10.1107/S0021889892001663. PubMed PMID:WOS:A1992JH63000005.

65. Franke D, Svergun DI. DAMMIF, a program for rapid ab-initio shape determination in small-angle scattering. J Appl Crystallogr. 2009;42(Pt 2):342–6. doi:10.1107/S0021889809000338. PubMed PMID:27630371; PubMed Central PMCID:PMCPMC5023043.

66. Volkov VV, Svergun DI. Uniqueness of ab initio shape determination in small-angle scattering. Journal of Applied Crystallography. 2003;36:860–4. doi:10.1107/S0021889803000268. PubMed PMID:WOS:000182284400105.

67. Petoukhov MV, Franke D, Shkumatov AV, Tria G, Kikhney AG, Gajda M, et al. New developments in the ATSAS program package for small-angle scattering data analysis. Journal of Applied Crystallography. 2012;45:342–50. doi:10.1107/S0021889812007662. PubMed PMID:WOS:000302808300025.

68. Kozin MB, Svergun DI. Automated matching of high- and low-resolution structural models. Journal of Applied Crystallography. 2001;34:33–41. doi:10.1107/S0021889800014126. PubMed PMID:WOS:000166532800007.

69. Yilmaz S, Drepper F, Hulstaert N, Cernic M, Gevaert K, Economou A, et al. Xilmass: A New Approach toward the Identification of Cross-Linked Peptides. Anal Chem. 2016;88(20):9949–57. doi:10.1021/acs.analchem.6b01585. PubMed PMID:27642655.

70. Yang B, Wu YJ, Zhu M, Fan SB, Lin J, Zhang K, et al. Identification of cross-linked peptides from complex samples. Nat Methods. 2012;9(9):904–6. doi:10.1038/nmeth.2099. PubMed PMID:22772728.

71. Chambers MC, Maclean B, Burke R, Amodei D, Ruderman DL, Neumann S, et al. A cross-platform toolkit for mass spectrometry and proteomics. Nat Biotechnol. 2012;30(10):918–20. doi:10.1038/nbt.2377. PubMed PMID: 23051804; PubMed Central PMCID:PMCPMC3471674.

72. Chan A, Schummer A, Fischer S, Schroter T, Cruz-Zaragoza LD, Bender J, et al. Pex17p-dependent assembly of Pex14p/Dyn2p-subcomplexes of the peroxisomal protein import machinery. Eur J Cell Biol. 2016;95(12):585–97. doi:10.1016/j.ejcb.2016.10.004. PubMed PMID:27823812.

73. Kostan J, Salzer U, Orlova A, Toro I, Hodnik V, Senju Y, et al. Direct interaction of actin filaments with F-BAR protein pacsin2. EMBO Rep. 2014;15(11):1154–62. doi:10.15252/embr.201439267. PubMed PMID:25216944; PubMed Central PMCID:PMCPMC4253489.

74. Marley J, Lu M, Bracken C. A method for efficient isotopic labeling of recombinant proteins. J Biomol NMR. 2001;20(1):71–5. PubMed PMID:11430757.

75. Kay LE, Nicholson LK, Delaglio F, Bax A, Torchia DA. Pulse Sequences for Removal of the Effects of Cross-Correlation between Dipolar and Chemical-Shift Anisotropy Relaxation Mechanism on the Measurement of Heteronuclear T1 and T2 Values in Proteins. J Magn Reson. 1992;97(2):359–75. doi:10.1016/0022-2364(92)90320-7. PubMed PMID:WOS:A1992HL67500012.

76. Palmer AG, Cavanagh J, Wright PE, Rance M. Sensitivity Improvement in Proton-Detected 2-Dimensional Heteronuclear Correlation Nmr-Spectroscopy. J Magn Reson. 1991;93(1):151–70. doi:10.1016/0022-2364(91)90036-S. PubMed PMID:WOS:A1991FL34200012.

77. Delaglio F, Grzesiek S, Vuister GW, Zhu G, Pfeifer J, Bax A. NMRPipe: a multidimensional spectral processing system based on UNIX pipes. J Biomol NMR. 1995;6(3):277–93. PubMed PMID: 8520220.

78. Svergun D, Barberato C, Koch MHJ. CRYSOL– a Program to Evaluate X-ray Solution Scattering of Biological Macromolecules from Atomic Coordinates. Journal of Applied Crystallography. 1995;28(6):768–73. doi:10.1107/s0021889895007047. PubMed PMID:WOS:A1995TP80300013.

79. Lyskov S, Gray JJ. The RosettaDock server for local protein-protein docking. Nucleic Acids Res. 2008;36(Web Server issue):W233–8. doi:10.1093/nar/gkn216. PubMed PMID: 18442991; PubMed Central PMCID:PMCPMC2447798.

80. Leber Y, Ruparelia AA, Kirfel G, van der Ven PF, Hoffmann B, Merkel R, et al. Filamin C is a highly dynamic protein associated with fast repair of myofibrillar microdamage. Hum Mol Genet. 2016;25(13):2776–88. doi:10.1093/hmg/ddw135. PubMed PMID:27206985.

81. Mlynek G, Lehner A, Neuhold J, Leeb S, Kostan J, Charnagalov A, et al. The Center for Optimized Structural Studies (COSS) platform for automation in cloning, expression, and purification of single proteins and protein-protein complexes. Amino Acids. 2014;46(6):1565–82. doi:10.1007/s00726-014-1699-x. PubMed PMID:24647677.

82. Mlynek G, Kostan J, Leeb S, Djinović-Carugo K. Tailored Suits Fit Better: Customized Protein Crystallization Screens. Crystal Growth & Design. 2019;20(2):984–94. doi:10.1021/acs.cgd.9b01328.

83. Jurrus E, Engel D, Star K, Monson K, Brandi J, Felberg LE, et al. Improvements to the APBS biomolecular solvation software suite. Protein Sci. 2018;27(1):112–28. doi:10.1002/pro.3280. PubMed PMID:28836357; PubMed Central PMCID:PMCPMC5734301.

84. Risi C, Eisner J, Belknap B, Heeley DH, White HD, Schroder GF, et al. Ca(2+)-induced movement of tropomyosin on native cardiac thin filaments revealed by cryoelectron microscopy. Proc Natl Acad Sci U S A. 2017;114(26):6782–7. doi:10.1073/pnas.1700868114. PubMed PMID:28607071; PubMed Central PMCID:PMCPMC5495243.

85. Haywood NJ, Wolny M, Rogers B, Trinh CH, Shuping Y, Edwards TA, et al. Hypertrophic cardiomyopathy mutations in the calponin-homology domain of ACTN2 affect actin binding and cardiomyocyte Z-disc incorporation. Biochem J. 2016;473(16):2485–93. doi:10.1042/BCJ20160421. PubMed PMID:27287556; PubMed Central PMCID:PMCPMC4980809.

