## Supplementary Figure Captions for "Molecular basis of F-actin regulation and sarcomere assembly *via* myotilin"

Figure S1. Related to Figure 1. Myotilin displays conformational ensemble in solution.

(**A**) Determination of molecular weight (MW) across the RI/RALLS elution peak collected in parallel to SEC‑SAXS mode. MW estimates are shown in red, right-angle laser light scattering (RALLS) trace in orange and refractive index (RI) trace in blue. The MW_RALLS_ (70 ± 7 kDa). MW estimates corresponding to elution time points prior to the main elution peak suggest a small presence of higher oligomeric species such as dimers. (**B**) Dimensionless Kratky plot indicates that while Trx-MYOT does contain ordered regions (corresponding to the Trx moiety and the Ig1 and Ig2 domains), most of the polypeptide chain is disordered and extended. In comparison, dimensionless Kratky plot of BSA, representing a typical folded protein, is shown. (**C, D**) D_max_ and R_g_ distributions from EOM analysis of Trx-MYOT. The parameters derived from the selected pool compared to the original pool suggest that Trx-MYOT is slightly restrictive in its flexibility and occupies a more compact conformation (EOM fit and models are shown in **Figures 1C and 1E**). (**E**) Schematic representation of Ig1Ig2^220-452^ depicting potential modes of dimerization.

(**F**) Theoretical *P(r)* *vs.* *r* plots calculated for potential Ig1Ig2^220-452^ dimerization modes showed in (**E**). Of these, both parallel and antiparallel dimers using tandem Ig1Ig2 as dimerization interface (parallel^(via Ig1Ig2)^, antiparallel^(via Ig1Ig2)^) display marginal increase in the D_max_. Staggered dimers display notable increase in the D_max_ similar to experimentally observed (**Figure 1F and Table S1**). In comparison to experimental data, where Ig1Ig2^220-452^ adopts a conformational ensemble in solution (**Figure 1E**) [1], *P(r)* *vs.* *r* plots for different dimerization modes were calculated using one (static) conformation, consequently resulting in a longer D_max_ compared to experimentally derived. In order to compare various *P(r)* functions, *P(r)* was normalized to the peak height. (**G**) *P(r)* *vs.* *r* plot for the concentration series of Ig1Ig2^220-452^ and Ig1Ig2^220-452 R405K^ measured at the same experimental conditions (for details see **Table S1**). Inset, respective concentration series with the corresponding SAXS profiles. Note, both Ig1Ig2^220-452^ and Ig1Ig2^220-452 R405K^, showed comparable concentration dependent increase in D_max_ suggesting that *in vitro* R405K mutation does not impair myotilin homo-dimerization.

Figure S2. Related to Figure 2. Myotilin binds to F-actin *via* Ig domains and disordered flanking regions and influences its dynamics.

(**A, B**) Binding of various myotilin constructs to F-actin in conditions B1 (**A**) or conditions B2 (**B**). The exponential binding curves fitted for each set of data points were used to determine apparent binding affinities shown in **Figure 2A**. Plotted values represent the mean ± SEM from 3-5 independent experiments. (**C**) Melting temperature (T_m_) of Trx-MYOT in various buffers as assessed by DSF assay (for details see Material and Methods). The pH of each buffer (blue rectangle) is indicated. T_m_ max (violet), denotes the buffer conditions in which T_m_ of Trx-MYOT was the highest (100 mM MES, pH 5.5). Brown line, indicates T_m_ in the standard buffer (100 mM Tris, pH 8.0). Note trend in increasing the stability of Trx-MYOT by lowering the pH. (**D**) Structural analysis of the functional (F-actin binding) sites on various Ig domains. Top panel: Basic amino acid residues with surface-exposed sidechains are shown as sticks (cyan). For palladin and filamin A this region corresponds to the F-actin binding region. Bottom panel: The functionally important residues as predicted by evolutionary coupling analysis and folding server EVfold [2] are shown as spheres, and coincide with the residues shown in the top panel on the lateral sides of the Ig domains. More intense (darker) color depicts a higher probability that the residue represents a functional site. (**E**) Binding of various myotilin constructs and control (GST) to fluorescently labelled monomeric DVD-actin measured by microscale thermophoresis (MST). (**F**) Table of the constructs and their affinity to DVD-actin obtained from the data shown in (**E**).

Figure S3. Related to Figure 3. Integrative model of myotilin:F-actin complex.

(**A-C**) Cross‐linking of myotilin Ig1Ig2^250-444^ (**A**), Ig1Ig2^185-454^ (**B**) and Ig1Ig2^185-498^ (**C**) with F-actin. Myotilin-actin complexes were cross-linked with DMTMM, either at increasing concentrations of DMTMM and fixed amount of proteins (Ig1Ig2^250-444^), or at its fixed concentration and varying amounts of proteins (Ig1Ig2^185-454^ and Ig1Ig2^250-498^). SDS-PAGEs of cross-linked samples stained with Coomassie Brilliant Blue are shown. Specific bands, corresponding to 1:1 complex were analyzed by MS and data obtained from it are summarized in **Table S2**. (**D**) Relative change of ^1^H-^15^N HSQC cross-peak intensities upon addition of F-actin to Ig1^250-344^ (blue, left panel) or Ig2^349-459^ (red, right panel). Values were averaged by a sliding window function over 11 amino acids. Stronger reduction in signal intensity of Ig2^349-459^ as compared to Ig1^250-344^ indicates that Ig2 binds more tightly to F-actin than Ig1. In all experiments, except for the one indicated by an asterisk (67 μM), a fixed concentration of myotilin was used (100 μM), while the concentration of F-actin was varied as indicated in the figure. (**E**) ^1^H-^15^N HSQC monitored shift changes in Ig1Ig2^250-444^ (50 µM) upon addition of F-actin (4 µM). Shifts were averaged by a sliding window function over 11 amino acids. In Ig1Ig2^250-444^ four main regions with the most pronounced effect upon addition of F-actin were identified (boxed green). Inset, topology diagram of I-type Ig domain. (**F**) Shifts used in (**E**) expressed using z-scores (see Methods for pertinent details on calculation of z-score). For all regions indicated in (**E**) we find more than one shift with a z-score above one and at least one higher than two. (**G**) Structural model of a single full-length myotilin molecule bound to F-actin, the residues identified in XL-MS analysis are highlighted in orange.

Figure S4. Related to Figure 4. Myotilin regulates binding of tropomyosin to F-actin and does not interact with PI(4,5)P_2_.

(**A**) Effects of myotilin on tropomyosin:F-actin interaction. Example of the SDS-PAGE used to generate **Figure 4B** is shown. Tropomyosin (Tpm) was incubated with F-actin before addition of Trx-MYOT at molar ratios indicated on the figure. F-actin and proteins bound were sedimented by centrifugation, and equal amounts of supernatant (S) and pellet (P) fractions were subjected to SDS-PAGE. (**B**) Effect of myotilin on α-actinin:F-actin interaction. Example of the SDS-PAGE used to generate **Figure 4D** is shown. α-Actinin-2 NEECK (ACTN2-NEECK) was incubated with F-actin before addition of tropomyosin (Tpm) at molar ratios indicated on the figure. F-actin and proteins bound were sedimented by centrifugation, and equal amounts of supernatant (S) and pellet (P) fractions were subjected to SDS-PAGE. (**C**) Model of tandem Ig domains of myotilin, actin binding domain of α-actinin-2 (ACTN2 ABD) and tropomyosin bound to F-actin was generated by superposition of our myotilin:tropomyosin:F-actin model shown on **Figure 4A** with the structures of F-actin bound to α-actinin-2, spectrin and filamin A, ABDs, at nominal resolutions of 16, 6.9 and 3.6 Å, respectively [3-5]. In the next step, crystal structure of α-actinin-2 ABD (salmon color, PDB: 5A36) [6] was superimposed over ABD of filamin A (homologous to ABD of cardiac and skeletal muscles expressed isoform filamin C) to obtain the final model. Dotted line (black) indicates position of tandem Ig domains of myotilin (myot Ig1, myot Ig2) on F-actin as shown on **Figure 4A**. N-terminal regions preceding the ABDs increase affinity to F-actin and are isoform specific [3]. The potential trace of the α-actinin-2 N-terminal extension (N-term.) on F-actin, as deduced from high resolution structures of F-actin decorated by filamin A and spectrin ABDs is shown with a dashed red line [5]. In addition, actin-binding sites of α-actinin-2 and myotilin clearly overlap, explaining mutually exclusive binding of myotilin, α-actinin-2 and tropomyosin to F-actin. Residues of F-actin found in cross‑links with N- and C-terminal regions flanking myotilin Ig domains are shown in orange (see **Figure 3A**). (**D** and **E**) Binding of myotilin to PI(4,5)P_2_. Tandem of myotilin Ig domains (Ig1Ig2^250-444^), Ig3 domain of palladin (Palladin Ig3) and double C2-like domain-containing protein beta (Doc2b) were incubated alone (control) or with different concentrations of PI(4,5)P_2_ (0–20%) in POPC vesicles and centrifuged. Supernatant (S) and pellet (P) fractions obtained were separated on SDS-PAGE. (**D**) Representative gels from one of the three independent experiments are shown. (**E**) Quantitative representation of (**D**), where the amount of protein found in pellet (protein in pellet) is represented as the percentage of the total amount of the protein found in both supernatant and pellet fractions. Mean values (± SEM) of three independent experiments are shown. Significance was assessed using 2‐tailed Student's *t*‐test, *p < 0.05 and **p < 0.01. Note, myotilin does not bind to PI(4,5)P_2_ containing vesicles. (**F**) Structural based sequence alignment of myotilin Ig1, myotilin Ig2 and palladin Ig3 domain. Residues that represent the putative PI(4,5)P_2_ and Ins(1,4,5)P_3_ binding sites on the Ig3 domain of palladin [7] are shown in green boxes. Conserved residues are shown in red. The numbers of the first and the last residue of the aligned sequences are given.
