## Supplementary figures and images for "Molecular basis of F-actin regulation and sarcomere assembly *via* myotilin"

### Supplementary Figures S1 - S4

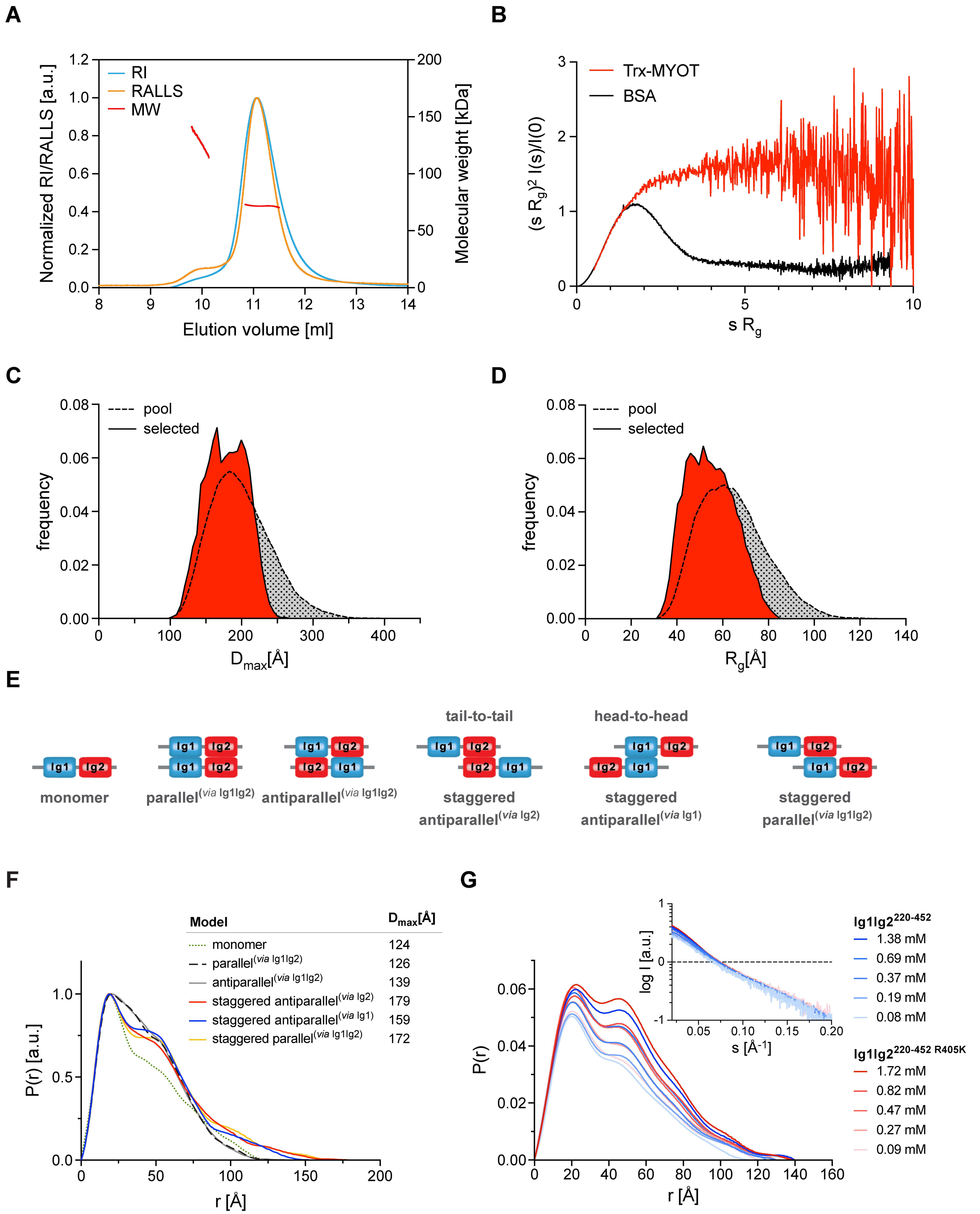

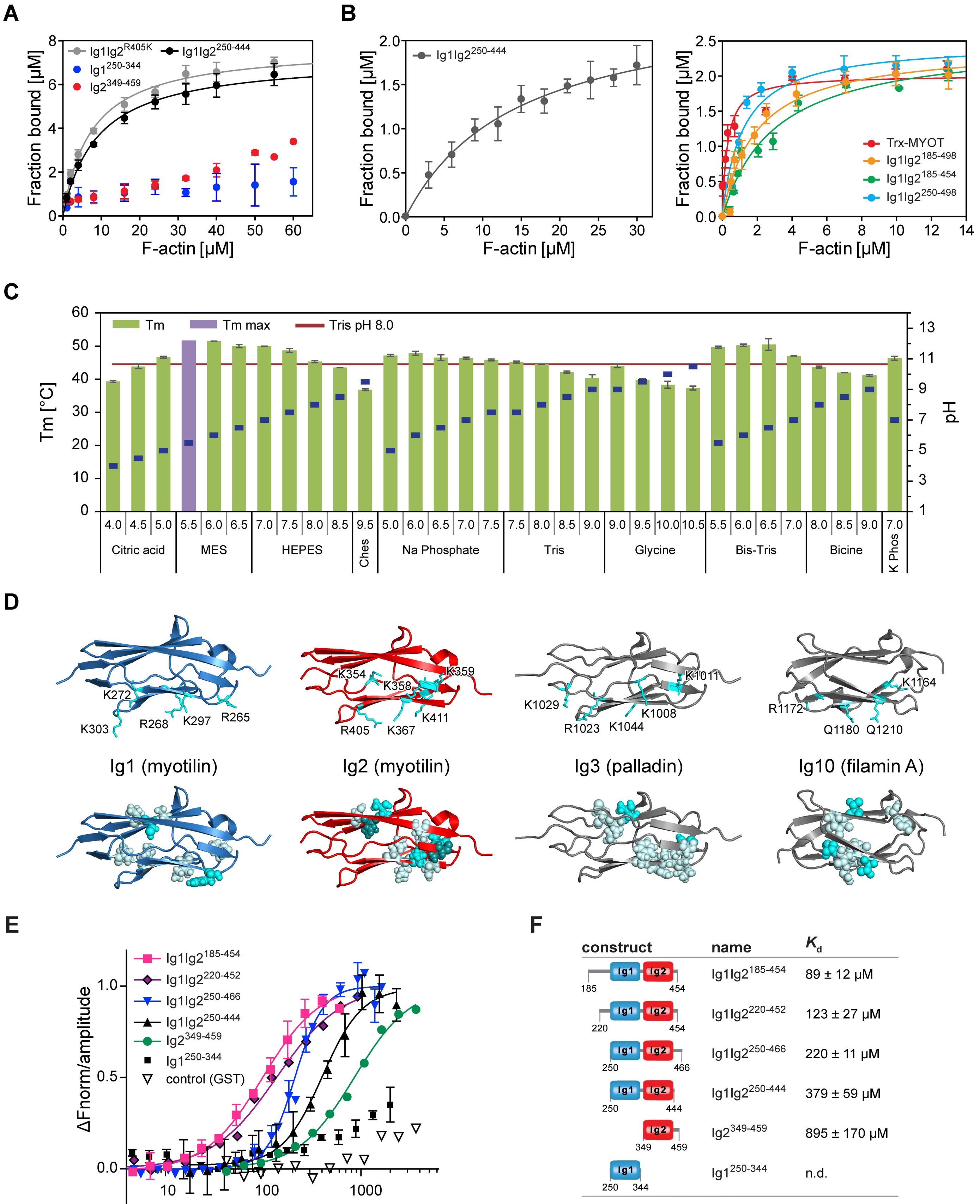

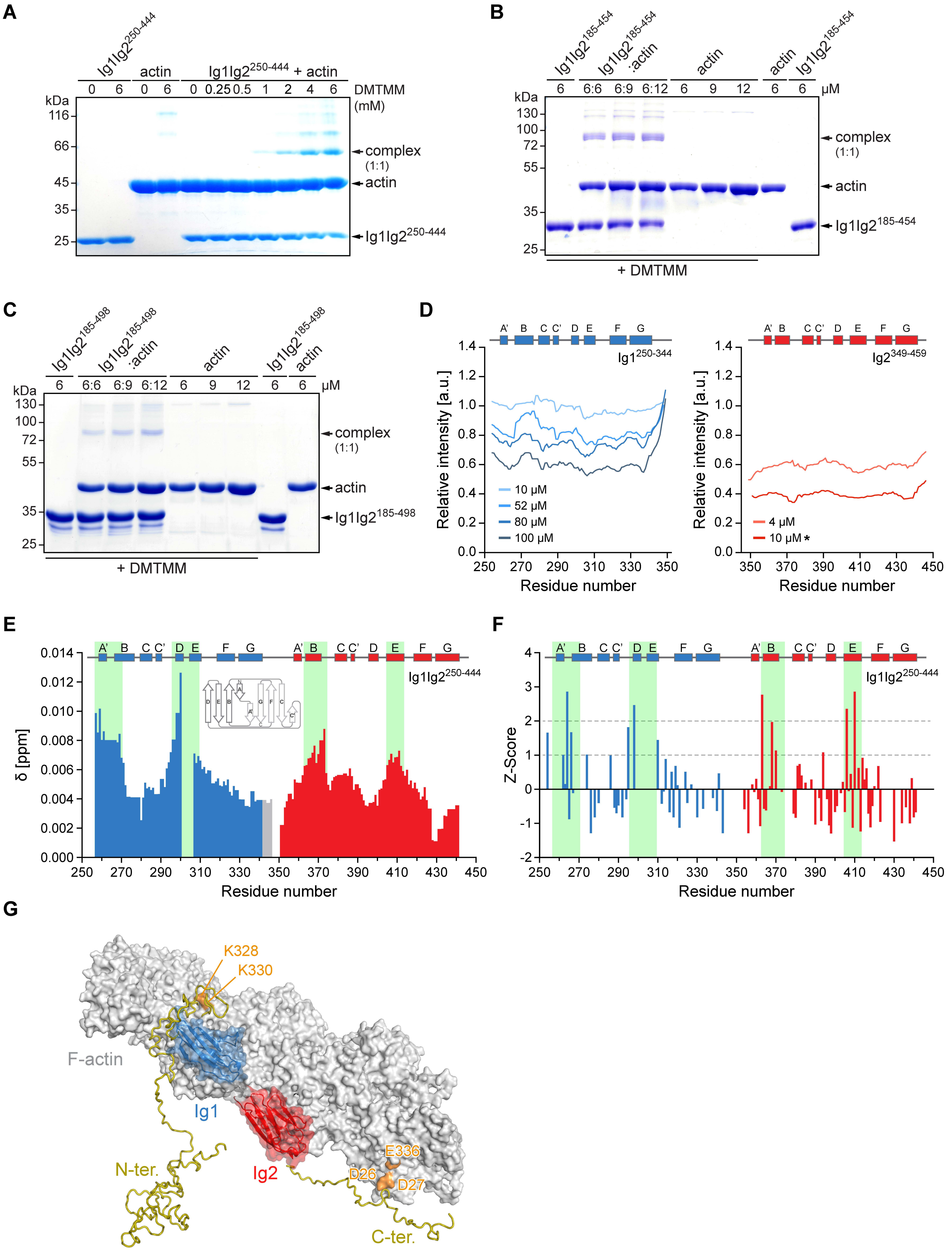

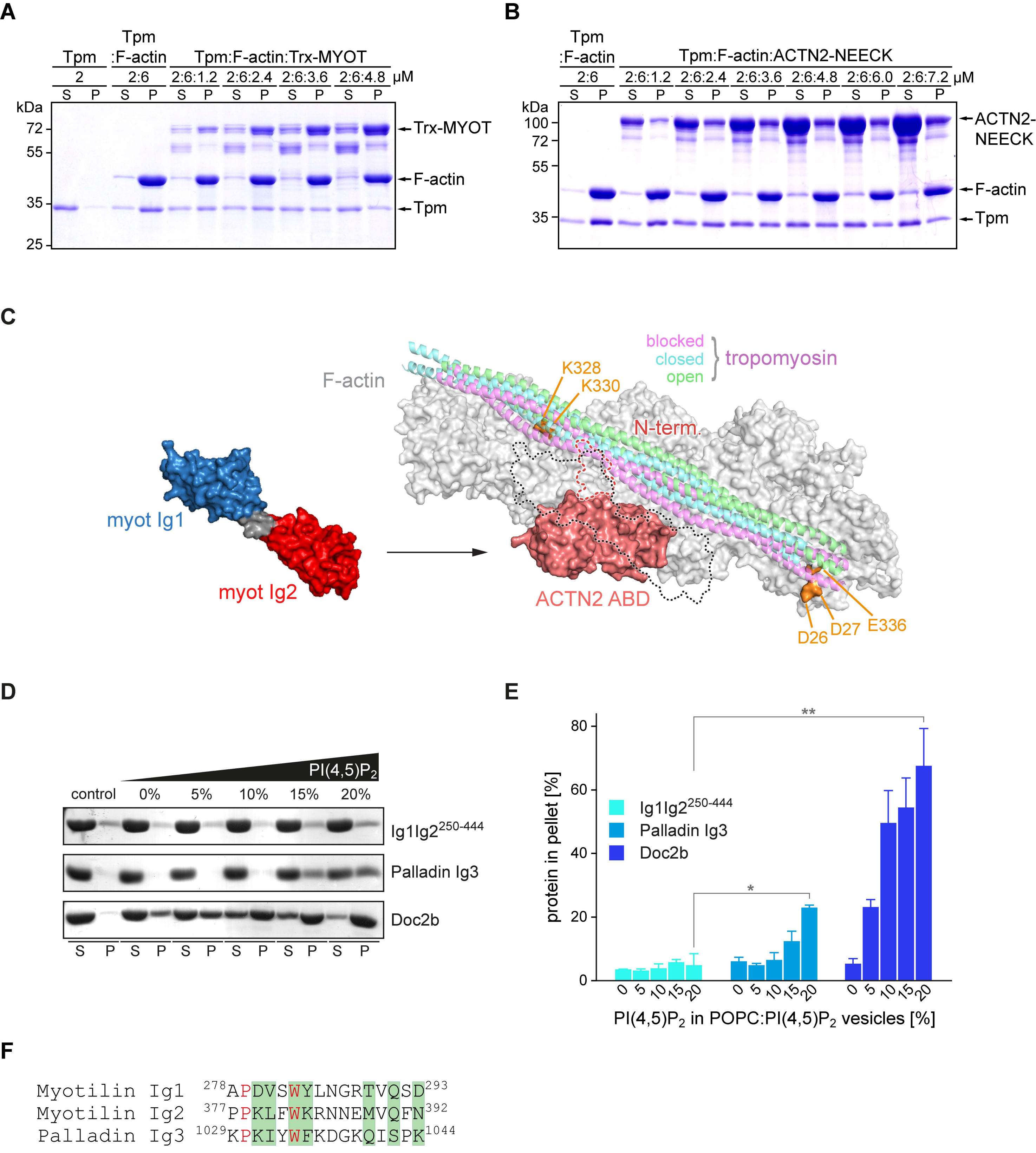
