## Supplementary Tables S1 - S3 for "Molecular basis of F-actin regulation and sarcomere assembly *via* myotilin"

### Table S1: SAXS data collection, analysis and derived structural parameters

| **Construct** | **Ig1Ig2$^{250-444}$*** | **Ig1Ig2$^{250-498}$** | **Ig1Ig2$^{220-452}$** | **Trx-MYOT** | **Ig1Ig2$^{220-452\dagger}$** | **Ig1Ig2$^{220-452\ R405K}$** |
|---|---|---|---|---|---|---|
| **Data collection parameters** | | | | | | |
| Radiation source | ESRF (Grenoble, France) | | DESY (Hamburg, Germany) | | | |
| | | | Doris | Petra III | | |
| Beamline | BM29 BioSAXS | | EMBL X33 | EMBL P12 | | |
| Detector | Pilatus 1M | | Pilatus 1M-W | Pilatus 2M | Pilatus 6M | |
| Beam geometry [mm, FWHM] | 0.10 × 0.20 | | n.d. | 0.12 × 0.20 | 0.12 × 0.20 | |
| Wavelength [nm] | 0.099 | | 0.126 | 0.124 | 0.124 | |
| Sample–detector distance [m] | 2.867 | | 2.7 | 3.1 | 3.0 | |
| Momentum of transfer *s* range [nm$^{-1}$] | 0.04–4.0 | | 0.1–6.0 | 0.08–3.5 | 0.15–7.3 | |
| Exposure time [s] | 15 | | 15 | 1 (SEC-SAXS) | 0.145 | |
| Temperature [°C] | 20 | | 11 | 20 | 20 | |
| Buffer | 20 mM HEPES, 150 mM NaCl, 5% glycerol, 1 mM DTT, pH 7.4 | | 20 mM MES, 200 mM NaCl, 3% glycerol, pH 6.0 | 20 mM Tris 400 mM NaCl, 250 mM arginine, 5% glycerol, pH 7.5 | 20 mM HEPES, 150 mM NaCl, 5% glycerol, 1 mM DTT, pH 7.4 | |
| **Overall parameters** | | | | | | |
| Conc. range measured$^{\$}$ | 1–14.12 mg/ml (0.05–0.64 mM) | 1–42.8 mg/ml (0.04–1.53 mM) | 1–52 mg/ml (0.04–1.96 mM) | n.a. | 2.1–36.6 mg/ml (0.08–1.38 mM) | 2.3–45.5 mg/ml (0.09–1.72 mM) |
| $R_g$ (Guinier) [Å] | 28 (0.64 mM) | 32 (0.54 mM)<br>37 (1.53 mM) | 31 (0.04 mM)<br>37 (0.79 mM)<br>39 (1.96 mM) | 51 | 30 (0.08 mM)<br>33 (0.19 mM)<br>33 (0.37 mM)<br>34 (0.69 mM)<br>36 (1.38 mM) | 33 (0.09 mM)<br>34 (0.27 mM)<br>34 (0.47 mM)<br>35 (0.82 mM)<br>36 (1.72 mM) |
| $R_g$ from PDDF [Å] | 29 (0.64 mM) | 34 (0.54 mM)<br>38 (1.53 mM) | 32 (0.04 mM)<br>39 (0.79 mM)<br>41 (1.96 mM) | 53 | 33 (0.08 mM)<br>34 (0.19 mM)<br>35 (0.37 mM)<br>37 (0.69 mM)<br>37 (1.38 mM) | 34 (0.09 mM)<br>36 (0.27 mM)<br>36 (0.47 mM)<br>37 (0.82 mM)<br>38 (1.72 mM) |
| $D_{max}$ [Å] | 101 (0.64 mM) | 140 (0.54 mM)<br>152 (1.53 mM) | 107 (0.04 mM)<br>151 (0.79 mM)<br>152 (1.96 mM) | 200 | 115 (0.08 mM)<br>125 (0.19 mM)<br>130 (0.37 mM)<br>140 (0.69 mM)<br>140 (1.38 mM) | 120 (0.09 mM)<br>130 (0.27 mM)<br>140 (0.47 mM)<br>140 (0.82 mM)<br>140 (1.72 mM) |
| $M_w$ (RALLS) [kDa] | n.a. | n.a. | n.a. | 70 | n.a. | n.a. |
| $M_w$ (DATMOV) [kDa] | 24.3 (0.64 mM) | 31.2 (0.54 mM)<br>44.7 (1.53 mM) | 27.3 (0.04 mM)<br>44.7 (0.79 mM)<br>58.1 (1.96 mM) | 63.7 | 30.8 (0.08 mM)<br>30.7 (0.19 mM)<br>33.7 (0.37 mM)<br>37.8 (0.69 mM)<br>43.2 (1.38 mM) | 29.1 (0.09 mM)<br>31.9 (0.27 mM)<br>35.6 (0.47 mM)<br>40.6 (0.82 mM)<br>45.7 (1.72 mM) |

*(continued)*

*(continued)*

| **Construct** | **Ig1Ig2$^{250\text{-}444*}$** | **Ig1Ig2$^{250\text{-}498}$** | **Ig1Ig2$^{220\text{-}452}$** | **Trx-MYOT** | **Ig1Ig2$^{220\text{-}452\dagger}$** | **Ig1Ig2$^{220\text{-}452\ R405K}$** |
|---|---|---|---|---|---|---|
| $M_w$ from Porod volume [kDa] | 21.4 (0.64 mM) | 26.4 (0.54 mM)<br>37.9 (1.53 mM) | 27.5 (0.04 mM)<br>36.7 (0.79 mM)<br>48.2 (1.96 mM) | 77.3 | 26.9 (0.08 mM)<br>25.9 (0.19 mM)<br>28.3 (0.37 mM)<br>30.9 (0.69 mM)<br>36.2 (1.38 mM) | 24.5 (0.09 mM)<br>26.3 (0.27 mM)<br>29.6 (0.47 mM)<br>33.6 (0.82 mM)<br>38.2 (1.72 mM) |
| $M_w$ (monomer sequence) [kDa]$^{\&}$ | 21.9 | 28.0 | 26.5 | 69.7 | 26.5 | 26.5 |
| **Software employed** | | | | | | |
| Primary data red. | SaxsAnalysis pipeline system | | | SASFLOW | SaxsAnalysis pipeline system | |
| Data processing | PRIMUS | | | PRIMUS / Cromixs | PRIMUS | |
| Calculation and comparison of scattering data | Crysol / Oligomer | | | | n. a. | |
| *Ab initio* modelling | DAMMIF | | | | n. a. | |
| Addition of missing residues | CORAL | | | | n. a. | |
| Ensemble modelling | EOM | | | | n. a. | |
| SASDB accession code | SASDF38 | SASDF48 | SASDF28 | SASDFZ7 | | |

*, the data for Ig1Ig2$^{250\text{-}444}$ were already measured and described previously (Puz et al., 2017)

$, for Ig1Ig2$^{250\text{-}498}$ and Ig1Ig2$^{220\text{-}452}$ structural parameters at various concentrations, relevant for the comparison, were calculated separately

&, $M_w$ (monomer sequence) denotes molecular weight calculated from the amino acid sequence of the monomeric species

†, measured at the same experimental settings/conditions as Ig1Ig2$^{220\text{-}452\ R405K}$

**Table S2: List of major cross-links found between myotilin and F-actin (using DMTMM)**

**Myotilin constructs used** **(A)** 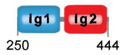

**(B)** 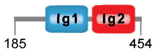

**(C)** 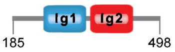


| From Protein | From Residue | To Protein | To Residue | Myotilin region | Myotilin fragment identified | Score |
|---|---|---|---|---|---|---|
| myotilin | **K 474** | actin | **E 102** | C-ter. | C | 212.6 |
| myotilin | **K 462** | actin | **D 27** | C-ter. | C | 199.5 |
| myotilin | **K 411** | actin | **D 27*** | Ig2 | A, B | A) 157.5, B) 175.5 |
| myotilin | **K 469** | actin | **E 102** | C-ter. | C | 169.4 |
| myotilin | **D 236** | actin | **K 330** | N-ter. | B, C | B) 138.4, C) 76.2 |
| myotilin | **D 236** | actin | **K 328*** | N-ter. | B | 134.9 |
| myotilin | **K 246** | actin | **E 336** | N-ter. | B, C | B) 105.9, C) 111.2 |
| myotilin | **E 245** | actin | **K 330** | N-ter. | B | 107.1 |
| myotilin | **K 246** | actin | **D 26** | N-ter. | C | 106.3 |
| myotilin | **K 452** | actin | **D 27*** | C-ter. | C | 103.9 |
| myotilin | **D 241** | actin | **K 330*** | N-ter. | B | 102.4 |
| myotilin | **K 469** | actin | **D 26*** | C-ter. | C | 102.0 |
| myotilin | **K 452** | actin | **D 26** | C-ter. | C | 100.9 |
| myotilin | **D 239** | actin | **K 330*** | N-ter. | B | 97.0 |
| myotilin | **D 239** | actin | **K 328** | N-ter. | B | 88.1 |
| myotilin | **K 474** | actin | **D 27** | C-ter. | C | 84.6 |
| myotilin | **K 474** | actin | **E 336*** | C-ter. | C | 82.7 |
| myotilin | **D 241** | actin | **K 328*** | N-ter. | B | 78.2 |
| myotilin | **K 462** | actin | **D 26*** | C-ter. | C | 77.7 |
| myotilin | **K 354** | actin | **D 27*** | Ig2 | C | 76.7 |
| myotilin | **K 246** | actin | **D 27*** | N-ter. | B, C | B) 54.5, C) 73.9 |
| myotilin | **K 415** | actin | **D 27** | Ig2 | A, B | A) 66.1, B) 57.6 |
| myotilin | **K 303** | actin | **D 27*** | Ig1 | A | 65.9 |
| myotilin | **K 367** | actin | **D 365** | Ig2 | B | 63.6 |
| myotilin | **K 367** | actin | **E 366** | Ig2 | B | 60.2 |
| myotilin | **K 452** | actin | **E 363** | C-ter. | C | 48.2 |
| myotilin | **K 411** | actin | **D 26** | Ig2 | A | 47.8 |
| myotilin | **K 474** | actin | **D 26** | C-ter. | C | 35.4 |

*, cross-links shown on Figure 3A; N-ter., region flanking Ig1Ig2 N-terminally; C-ter., region flanking Ig1Ig2 C-terminally; Score, computed as -10 log10 (E-value) representing best identification for a crosslink pair. The higher values of the score indicate more reliable identifications.

### Table S3: List of constructs

| Name | [1]Protein | Residues | Mutations | Vector | [2]Tag | Expression in | Origin |
|---|---|---|---|---|---|---|---|
| **Trx-MYOT** | myotilin | 1-498 | - | pETM-20 | N-Trx-$His_6$-TEV | *E. coli* | this study |
| **Trx-MYOT-NEECK** | myotilin | 1-498 | F96E, L97E, L101E | pETM-20 | N-Trx-$His_6$-TEV | *E. coli* | this study |
| **Ig1Ig2$^{185-498}$** | myotilin | 185-498 | - | pETM-20 | N-Trx-$His_6$-3C C-Strep | *E. coli* | this study |
| **[3]Ig1Ig2$^{185-498}$*** | myotilin | 185-498 | - | pDB-HisGST | N-$His_6$-GST-TEV | *E. coli* | this study |
| **Ig1Ig2$^{220-452}$** | myotilin | 220-452 | - | pETM-14 | N-$His_6$-3C | *E. coli* | this study |
| **Ig1Ig2$^{220-452\ R405K}$** | myotilin | 220-452 | R405K | pETM-14 | N-$His_6$-3C | *E. coli* | this study |
| **[4]Ig1Ig2$^{250-498}$** | myotilin | 250-498 | - | pETM-14 | N-$His_6$-3C | *E. coli* | this study |
| **Ig1Ig2$^{250-444}$** | myotilin | 250-444 | - | pETM-14 | N-$His_6$-3C | *E. coli* | Puz et al., 2017 |
| **Ig1Ig2$^{K354A}$** | myotilin | 250-444 | K354A | pETM-14 | N-$His_6$-3C | *E. coli* | this study |
| **Ig1Ig2$^{Q356A}$** | myotilin | 250-444 | Q356A | pETM-14 | N-$His_6$-3C | *E. coli* | this study |
| **Ig1Ig2$^{K358A}$** | myotilin | 250-444 | K358A | pETM-14 | N-$His_6$-3C | *E. coli* | this study |
| **Ig1Ig2$^{K359A}$** | myotilin | 250-444 | K359A | pETM-14 | N-$His_6$-3C | *E. coli* | this study |
| **Ig1Ig2$^{K367A}$** | myotilin | 250-444 | K367A | pETM-14 | N-$His_6$-3C | *E. coli* | this study |
| **Ig1Ig2$^{R405K}$** | myotilin | 250-444 | R405K | pETM-14 | N-$His_6$-3C | *E. coli* | this study |
| **Ig1Ig2$^{K411A}$** | myotilin | 250-444 | K411A | pETM-14 | N-$His_6$-3C | *E. coli* | this study |
| **Ig1Ig2$^{K354/359A}$** | myotilin | 250-444 | K354A, K359A | pETM-14 | N-$His_6$-3C | *E. coli* | this study |
| **Ig1Ig2$^{K354/358/359A}$** | myotilin | 250-444 | K354A, K358A, K359A | pETM-14 | N-$His_6$-3C | *E. coli* | this study |
| **Ig1$^{250-344}$** | myotilin | 250-344 | - | pET-3d(+) | N-$His_6$-TEV | *E. coli* | this study |
| **Ig2$^{349-459}$** | myotilin | 349-459 | - | pET-3d(+) | N-$His_6$-TEV | *E. coli* | this study |
| **MYOT$^{WT}$** | myotilin | 1-498 | - | pEGFP-N1 | N-EGFP | C2C12 | this study |
| **MYOT$^{K354A}$** | myotilin | 1-498 | K354A | pEGFP-N1 | N-EGFP | C2C12 | this study |
| **MYOT$^{K359A}$** | myotilin | 1-498 | K359A | pEGFP-N1 | N-EGFP | C2C12 | this study |
| **MYOT$^{K354/359A}$** | myotilin | 1-498 | K354A, K359A | pEGFP-N1 | N-EGFP | C2C12 | this study |
| **MYOT$^{K354/358/359A}$** | myotilin | 1-498 | K354A, K358A, K359A | pEGFP-N1 | N-EGFP | C2C12 | this study |
| **ACTN2-WT** | α-actinin-2 | 1-894 | - | pET-3d(+) | N-$His_6$-TEV | *E. coli* | Ribeiro et al., 2014 |
| **ACTN2-NEECK** | α-actinin-2 | 1-894 | R268E, I269E, L273E | pET-3d(+) | N-$His_6$-TEV | *E. coli* | Ribeiro et al., 2014 |
| **ACTN2-EF14** | α-actinin-2 | 746-894 | - | pETM-14 | N-$His_6$-3C | *E. coli* | this study |
| **palladin Ig3** | palladin | 1022-1126 | - | pTBSG | N-$His_6$-TEV | *E. coli* | Yadav et al., 2016 |
| **Doc2b** | Doc2b | 125-412 | - | pGEX4T1 | N-GST-Thr | *E. coli* | Groffen et al., 2010 |
| **DVD-actin** | actin-5C | 1-376 | D287A, V288A, D289A | pFastBacHT | N-$His_6$ | *Sf9* | Zahm et al., 2013 |
| **tropomyosin** | tropomyosin | | | | | *E. coli* | von der Ecken et al., 2015 |

(1), myotilin (human, Uniprot-ID Q9UBF9); α-actinin-2 (human, P35609), palladin (Mus musculus, Q9ET54), Doc2b (Rattus norvegicus, P70610), actin-5C (Drosophila melanogaster, P10987), tropomyosin (Tpm1.1st, Homo sapiens, NP_001018005.1)

(2), in many cases removed during purification; N, N-terminal fusion; Trx, Thioredoxin-tag; $His_6$, 6xHis-tag, TEV; TEV protease cleavage site; 3C, HRV-3C protease cleavage site; C, C-terminal fusion; Strep, Strep-tag II; Thr, thrombin cleavage site, GST, Glutathione s-transferase-tag; EGFP, Enhanced green fluorescent protein-tag

(3), Ig1Ig2$^{185-498}$* was used to produce Ig1Ig2$^{185-454}$, and GST

(4), Ig1Ig2$^{250-498}$ was used to produce Ig1Ig2$^{250-466}$
